# Single-cell profiling of transcriptome and histone modifications with EpiDamID

**DOI:** 10.1101/2021.10.26.465688

**Authors:** Franka J. Rang, Kim L. de Luca, Sandra S. de Vries, Christian Valdes-Quezada, Ellen Boele, Phong D. Nguyen, Isabel Guerreiro, Yuko Sato, Hiroshi Kimura, Jeroen Bakkers, Jop Kind

## Abstract

Recent advances in single-cell sequencing technologies have enabled simultaneous measurement of multiple cellular modalities, including various combinations of transcriptome, genome and epigenome. However, comprehensive profiling of the histone post-translational modifications that influence gene expression at single-cell resolution has remained limited. Here, we introduce EpiDamID, an experimental approach to target a diverse set of chromatin types by leveraging the binding specificities of genetically engineered proteins. By fusing Dam to single-chain variable fragment antibodies, engineered chromatin reader domains, or endogenous chromatin-binding proteins, we render the DamID technology and all its implementations compatible with the genome-wide identification of histone post-translational modifications. Importantly, this enables the joint analysis of chromatin marks and transcriptome in a variety of biological systems at the single-cell level. In this study, we use EpiDamID to profile single-cell Polycomb occupancy in mouse embryoid bodies and provide evidence for hierarchical gene regulatory networks. We further demonstrate the applicability of this method to *in vivo* systems by mapping H3K9me3 in early zebrafish embryogenesis, and detect striking heterochromatic regions specifically in the notochord. Overall, EpiDamID is a new addition to a vast existing toolbox for obtaining systematic insights into the role of chromatin states during dynamic cellular processes.

## Introduction

Histone post-translational modifications (PTMs) are an important aspect of chromatin structure and gene regulation. The addition of these chemical groups to histone tails can modulate the accessibility to the underlying DNA and form a binding platform for myriad downstream effector proteins. Amongst others, this can result in the shielding or recruitment of transcription factors (TFs) to promoters and enhancers. As such, histone PTMs play key roles in a multitude of biological processes, including lineage specification (e.g., Juan et al., 2016; Nicetto et al., 2019; Pengelly et al., 2013), cell cycle regulation (e.g., Hirota et al., 2005; W. Liu et al., 2010), and response to DNA damage (e.g., Rogakou et al., 1998; Sanders et al., 2004).

Over the past decade, antibody-based DNA-sequencing methods, such as chromatin immunoprecipitation followed by sequencing (ChIP-seq), Cleavage Under Target and Release Under Nuclease (CUT&RUN) (Skene & Henikoff, 2017), and related techniques (Schmid et al., 2004), have provided valuable insights into the function of histone PTMs in a variety of biological contexts. However, the general requirement of high numbers of input cells consequently provides a population-average view of the assayed histone PTM that belies the complexity of many biological systems. As a response, several low-input methods have been developed that can assay histone PTMs in individual cells (Ai et al., 2019; Hainer et al., 2019; Harada et al., 2019; Ku et al., 2019; Rotem et al., 2015; Zeller et al., 2021). While these single-cell methods offer a first understanding of the epigenetic heterogeneity between cells, it remains challenging to establish a direct link to other measurable outputs such as transcription or cellular state. Recently, a variety of single-cell multi-modal techniques have been developed that can simultaneously probe one or multiple aspects of gene regulation in conjunction with transcription in individual cells (Angermueller et al., 2016; Argelaguet et al., 2019; J. Cao et al., 2018; Clark et al., 2018; Moudgil et al., 2020; Rooijers et al., 2019; Xiong et al., 2021; Zhu et al., 2019, 2021). These techniques thus provide a way to link gene regulatory mechanisms to transcriptional output and cellular state in an unprecedented manner.

We recently developed scDam&T-seq, a method that can assay both transcription and DNA-protein contacts in single cells by combining single-cell DamID and CEL-Seq2 (Rooijers et al., 2019). DamID-based techniques attain specificity by tagging a protein of interest (POI) with the *E. coli* Dam methyltransferase, which will methylate adenines in a GATC motif in the proximity of the POI (Filion et al., 2010; Vogel et al., 2007). DamID is especially suited for obtaining information from individual cells, because DNA-protein contacts are recorded directly on the DNA in the living cell, and sample processing is particularly efficient with little loss of material. However, since Dam cannot be tagged directly to a post-translationally modified proteins by genetic engineering, this has precluded the use of any DamID methods for studying these epigenetic marks.

Here, we present EpiDamID, an extension of existing DamID-based protocols for the study of histone PTMs that can be applied in single cells. In EpiDamID, Dam is fused to a targeting domain with specific affinity for the histone PTM of interest. These targeting domains can be either a) full-length proteins with endogenous binding affinity, b) protein domains from known chromatin binders (Kungulovski et al., 2014, 2016; Vermeulen et al., 2007), or c) *m*odification-specific *int*racellular anti*bodies* (mintbodies) (Sato et al., 2013, 2016; Tjalsma et al., 2021) (Fig. 1A). Since this approach is an adaptation that can be applied to any DamID protocol, it provides all advantages that the DamID toolbox has to offer and makes them available to the study of chromatin modifications. This includes the possibility to perform (live) imaging of Dam-methylated DNA (Altemose et al., 2020; Borsos et al., 2019; Kind et al., 2013), tissue-specific study of model organisms without cell isolation via Targeted DamID (TaDa) (Southall et al., 2013), DamID-directed proteomics (Wong et al., 2021), multi-modal single-molecule sequencing (Cheetham et al., 2021), (multi-modal) single-cell sequencing studies (Altemose et al., 2020; Borsos et al., 2019; Kind et al., 2015; Rooijers et al., 2019), and the processing of samples with extremely little starting material (Borsos et al., 2019).

**Figure 1.**
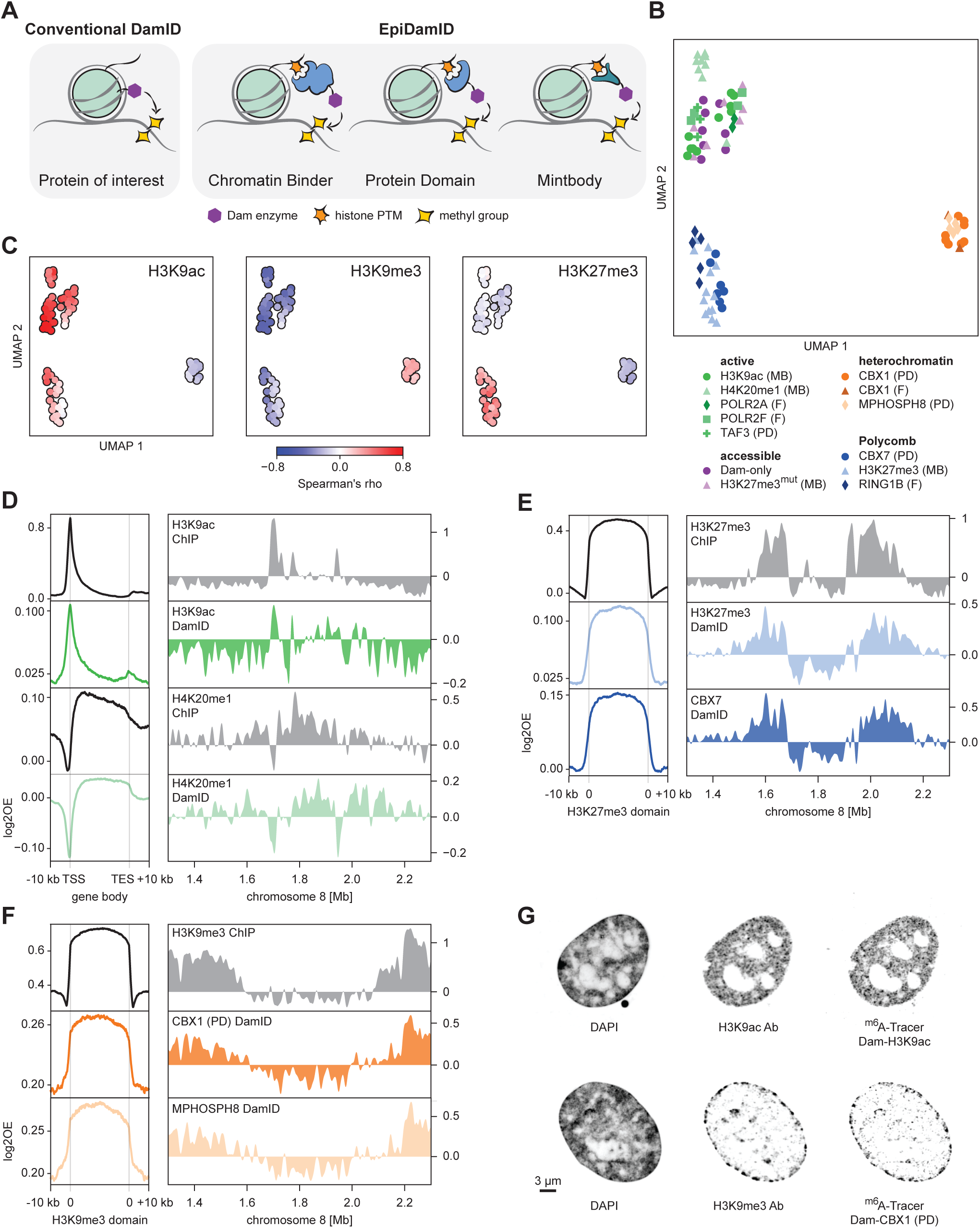
Targeting domains specific to histone modifications mark distinct chromatin types with EpiDamID. **A**, Schematic overview of EpiDamID concept compared to conventional DamID. **B**, UMAP of DamID samples, colored by targeting construct. The abbreviations MB, PD and F indicate what type of EpiDamID construct is used. MB: mintbody; PD: protein domain; F: full protein. **C**, UMAPs as in A, colored by correlation with selected ChIP-seq samples. **D-F**, Left, average DamID and ChIP-seq enrichment plots over genomic regions of interest. Signal (log2OE) is normalized for untethered Dam or input, respectively. Regions are gene bodies for H3K9ac and H4K20me1 (**D**), and ChIP-seq domains for H3K9me3 (**E**) and H3K27me3 (**F**). **D-F**, Right, genome browser view of ChIP-seq and DamID enrichment (log2OE) corresponding to left panels. The data shown in D-F represents the combined data of all samples of each targeting domain. **G**, Confocal images of nuclear chromatin showing DAPI, immunofluorescent staining against an endogenous histone modification, and its corresponding EpiDamID construct visualized with m6A-Tracer. Top: H3K9ac, bottom: H3K9me3.

We first validated the specificity of EpiDamID by targeting many different chromatin types in populations of human RPE-1 cells. To demonstrate the potential of EpiDamID, we then implemented the previously developed scDam&T-seq method (Rooijers et al., 2019) in mouse embryonic stem cells (mESCs) and obtained high-quality single-cell histone PTM profiles for selected targeting constructs. Next, we leveraged this single-cell resolution to study the Polycomb mark H3K27me3 and its relationship to transcription in mouse embryoid bodies (EBs), an in vitro differentiation system that mimics aspects of embryonic development (Desbaillets et al., 2000). We identified distinct Polycomb-regulated and Polycomb-independent hierarchical TF networks covering both lineage-specific and ubiquitous functions. Finally, we developed a protocol to assay cell type-specific patterns of the heterochromatic mark H3K9me3 in the zebrafish embryo and discovered broad domains of heterochromatin specific to the notochord. Together, these results show that EpiDamID provides a versatile tool that can be easily implemented in diverse biological settings to obtain histone PTM profiles of individual cells.

## Results

### Targeting domains specific to histone modifications mark distinct chromatin types with EpiDamID

The conventional DamID approach involves genetically engineering a POI to the bacterial methyltransferase Dam (Fig. 1A). In this study, we adapted the DamID method to detect histone PTMs by fusing Dam to one of the following: 1) full-length chromatin proteins, 2) tuples of well-characterized reader domains (Kungulovski et al., 2014, 2016; Vermeulen et al., 2007), or 3) single-chain variable fragments (scFv) also known as mintbodies (Sato et al., 2013, 2016; Tjalsma et al., 2021) (Fig. 1A). Such constructs have been previously successfully applied in microscopy, proteomics and ChIP experiments (Sato et al., 2013, 2016, 2021; Tjalsma et al., 2021; Villaseñor et al., 2020). The different constructs were categorized based on their targets into the following chromatin types: accessible, active, heterochromatin, and Polycomb. This approach is henceforth referred to as EpiDamID, and the construct fused to Dam as the targeting domain. We generated various expression constructs for each of the different targeting domains, testing promoters (HSP, PGK), orientations (Dam-POI, POI-Dam) and two versions of the Dam protein (Dam, Dam126) (Supplementary Table 1). The choice of promoter influences the expression level of the Dam-POI, whereas the orientation may affect access of the POI to its target. In the Dam126 mutant, the N126A substitution reduces its binding affinity to the DNA and consequently diminishes off-target methylation (Park et al., 2018; Szczesnik et al., 2019). We introduced the Dam constructs by viral transduction in hTERT-immortalized RPE-1 cells and performed DamID2 followed by high-throughput sequencing (Markodimitraki et al., 2020). To validate our data with an orthogonal method, we generated antibody ChIP-seq of various histone modifications.

The DamID samples were filtered on sequencing depth and information content (IC), a measure of the amount of signal in the data compared to the background genome-wide distribution of mappable fragments (Methods). IC is a valuable metric for determining overall sample quality and signal-to-noise levels (Fig. S1A-B). The IC additionally showed that tuples of reader domains fused to Dam perform better than single domains, in agreement with a recent study employing similar domains for proteomics purposes (Villaseñor et al., 2020) (Fig. S1A-B). Therefore, only data from the triple reader domains were included in further analyses.

Visualization of all filtered DamID samples by uniform manifold approximation and projection (UMAP) shows that EpiDamID mapping allows for identification of distinct chromatin types (Fig. 1B). Genome-wide DamID signal correlates well with antibody ChIP-seq signal of the same chromatin target (Fig. 1C and S1C). Importantly, DamID samples do not group based on construct type, promoter, Dam type, sequencing depth, or IC (Fig. S1D-E), indicating that those properties are separate from target specificity. To further compare DamID with ChIP-seq, we calculated enrichment over relevant genomic regions (genes or ChIP-seq peaks/domains). Cumulative signal shows excellent concordance between the methods and displays the expected patterns for all targets (Fig. 1D-F, left), as do example regions along the linear genome (Fig. 1D-F, right). It was previously reported that use of the Dam126 mutant improves signal quality compared to the use of wildtype (WT) Dam (Szczesnik et al., 2019). Indeed, we observed markedly improved sensitivity and reduced background methylation with the mutant Dam126 compared to WT Dam in our data (Fig. S1F-G).

Finally, we further validated the correct nuclear localization of Dam-marked chromatin with microscopy by immunofluorescent staining of endogenous histone PTMs and DamID visualization using ^m6^A-Tracer protein (Kind et al., 2013; Schaik et al., 2020) (Fig. 1G).

Together, these results show that EpiDamID specifically targets histone PTMs and enables identification of their genomic distributions by next-generation sequencing.

### Detection of histone PTMs in single mouse embryonic stem cells with EpiDamID

We next sought to establish whether the EpiDamID approach could be used to achieve single-cell profiles. To this end, we generated clonal, inducible mESC lines for each of the following targeting domains fused to Dam: H4K20me1 mintbody, H3K27me3 mintbody, and the H3K27me3-specific CBX7 protein domain (3x tuple). While H4K20me1 is enriched over the gene body of active genes (Shoaib et al., 2021), the heterochromatic mark H3K27me3 is enriched over the promoter of developmentally regulated genes until the appropriate moment of their activation during differentiation (Boyer et al., 2006; Riising et al., 2014). As controls, we included an H3K27me3^mut^ mintbody construct whose antigen-binding ability is abrogated via a point mutation in the third complementarity determining region of the heavy chain (Tyr105 to Phe), and a published mESC line expressing untethered Dam (Rooijers et al., 2019). Using these cell lines, we followed the scDam&T-seq protocol to generate 442 to 1,402 single-cell samples per construct. After filtering on the number of unique GATCs and IC, we retained 283 to 855 samples with high-quality DamID signal (Fig S2A-C). For the subsequent analyses, we also included a published dataset of Dam fused to RING1B (Rooijers et al., 2019) as an example of a full-length chromatin reader targeting Polycomb chromatin with DamID. All of these constructs contained the WT Dam, as we found that the reduced activity of Dam126 was insufficient to produce high-quality single-cell signal (data not shown).

Dimensionality reduction on the resulting single-cell datasets revealed that the samples primarily separated based on chromatin type (Fig. 2A), indicating that the various targeting domains result in specific methylation. To further confirm the specificity of the constructs, we used mESC H3K27me3 (ENCSR059MBO) and H3K9ac (ENCSR000CGP) ChIP-seq datasets from the ENCODE portal (Davis et al., 2018) and generated our own mESC H4K20me1 ChIP-seq dataset. For all single cells, we computed the enrichment of single-cell counts within H3K27me3, H3K9ac and H4K20me1 ChIP-seq domains. These results show a strong enrichment of EpiDamID counts within domains for the corresponding histone PTMs (Fig. 2B-D), indicating that the methylation patterns are specific for their respective chromatin targets, even at the single-cell level. We further validated the approach by combining single-cell samples per construct to obtain *in silico* population data, and computed the enrichment over H3K27me3 ChIP-seq domains (Fig. 2E) and active gene bodies (Fig. 2F) for the Polycomb-targeting constructs and H4K20me1, respectively. This illustrates that the combined signal, as well as the signal of the best single-cell samples, is strongly enriched over genomic regions of the corresponding histone PTM. Contrary to the H3K27me3 construct, its mutated mintbody control, H3K27me3^mut^, shows little enrichment over H3K27me3 ChIP-seq domains (Fig. 2B and Fig. S2D) further corroborating the specificity of the EpiDamID approach. Besides the average enrichment patterns, the specificity of the signal is also observed at individual loci in both the *in silico* populations and single cells (Fig. 2G-H and Fig. S2E).

**Figure 2:**
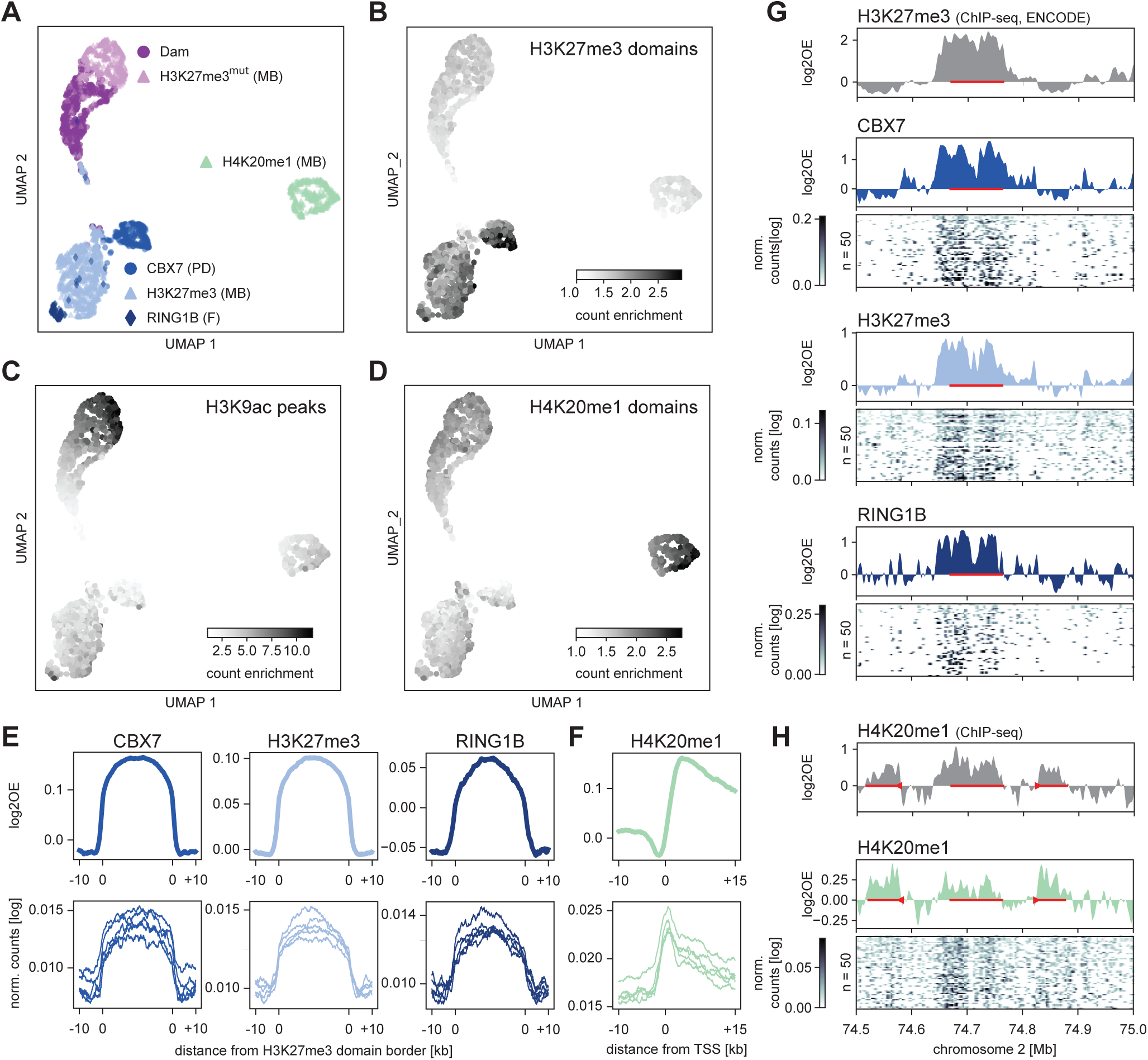
Detection of histone PTMs in single mouse embryonic stem cells with EpiDamID. **A,** UMAP based on the single-cell DamID readout of all single-cell samples. Samples are colored according to the targeting domain. The abbreviations MB, PD and F indicate what type of EpiDamID construct is used. MB: mintbody; PD: protein domain; F: full protein. **B-D**, DamID UMAP as in A, colored by the enrichment of counts within H3K27me3 ChIP-seq domains (B), H3K9ac ChIP-seq peaks (C), and H4K20me1 ChIP-seq domains (D). Count enrichment was computed as the fraction of GATC counts that fell within the regions, relative to the total fraction of genomic GATC positions inside these domains. **E**, Average signal over H3K27me3 ChIP-seq domains of CBX7 and H3K27me3 targeting domains and full-length RINGB1B protein. **F**, Average signal over the TSS of the top quartile active genes (as measured by H3K9ac ChIP-seq signal) of the H4K20me1 targeting domain. **E-F**, Top: *in silico* populations normalized for Dam; Bottom: five of the best single-cell samples (bottom) normalized only by read depth. **G-H**, Signal of various marks over the *HoxD* cluster and neighboring regions. ChIP-seq data is normalized for input control. The DamID tracks show the Dam-normalized *in silico* populations of the various Dam-fusion proteins, while heatmaps show the depth-normalized single-cell data of the fifty richest cells. The red bar around 74.7 Mb indicates the HoxD cluster. In **H**, the left red bar indicates the *Lnp* gene, the right bar indicates the *Mtx2* gene.

These results collectively demonstrate that both mintbodies and protein domains can be used to map histone PTMs in single cells with the EpiDamID approach.

### Joint profiling of Polycomb chromatin and gene expression in mouse embryoid bodies

To exploit the benefits of simultaneously measuring histone PTMs and transcriptome, and to test the potential of the method to capture chromatin dynamics, we chose to profile Polycomb in mouse EBs. We targeted the two main Polycomb repressive complexes (PRC) with EpiDamID using the full-length protein RING1B and H3K27me3-mintbody fused to Dam. RING1B is a core PRC1 protein that mediates H2AK119 ubiquitylation (de Napoles et al., 2004; H. Wang et al., 2004), and H3K27me3 is the histone PTM deposited by PRC2 (R. Cao et al., 2002; Czermin et al., 2002; Kuzmichev et al., 2002; Müller et al., 2002). Both PRC1 and PRC2 have key roles in gene regulation during stem cell differentiation and early embryonic development (see (Piunti & Shilatifard, 2021) and (Blackledge & Klose, 2021) for recent reviews on this topic).

To assay a diversity of cell types at different time points, we harvested EBs for scDam&T-seq at day 7, 10 and 14 post aggregation, next to ESCs grown in 2i/LIF (Fig. 3A). In addition to RING1B and H3K27me3-mintbody, we included the untethered Dam protein for all time points as a control for chromatin accessibility. Collectively, we obtained 2,943 cells that passed both DamID and transcriptome thresholds (Fig. S3A). Based on the transcriptional readout, we identified eight distinct clusters across all time points (Fig. 3B). We next integrated the EB transcriptome data with the publicly available mouse embryo atlas (Pijuan-Sala et al., 2019) to confirm the correspondence of the EB cell types with early mouse development, and guide cluster annotations (Fig. S3B-C). The results indicated the presence of both pluripotent and more differentiated cellular states, including epiblast, endoderm, and mesoderm lineages. Notably, the DamID readout alone was sufficient to consistently separate cells on chromatin type (Fig. 3C) and to distinguish between the pluripotent and more lineage-committed cells (Fig. 3D-E). Thus, the chromatin profiles in individual cells display cell type-specific patterns of chromatin accessibility and Polycomb association. Prompted by this observation, we trained a linear discriminant analysis (LDA) classifier to assign an additional 1,543 cells with poor transcriptional data to cell type clusters, based on their DamID signal (Fig. S3D-E).

**Figure 3.**
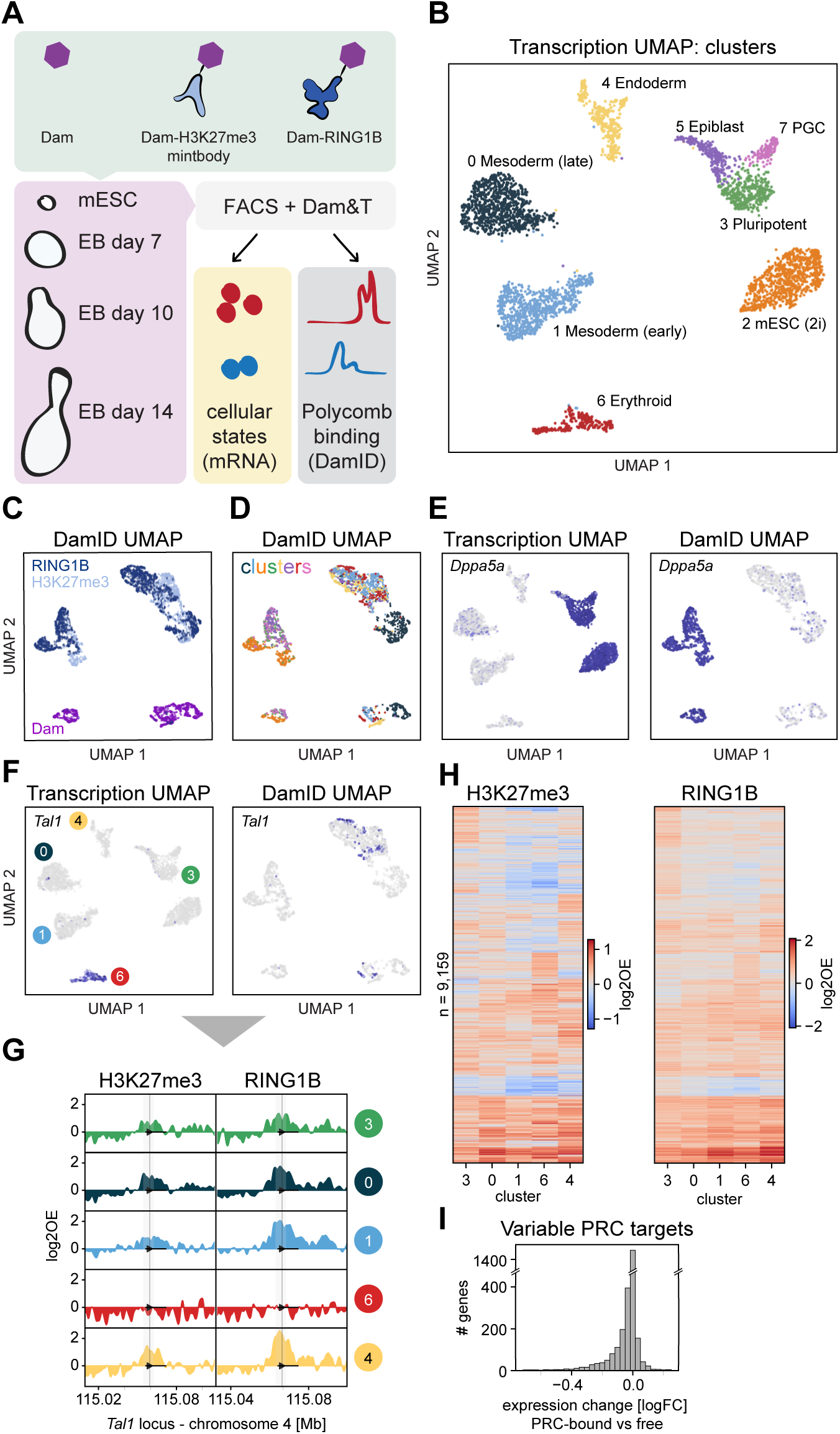
Joint profiling of Polycomb chromatin and gene expression in mouse embryoid bodies. **A,** Schematic showing the experimental design. **B,** UMAP of samples based on CEL-Seq2 readout, colored by cluster. **C-D,** UMAP of samples based on DamID readout, colored by construct (**C**) and cluster (**D**). **E**, Transcriptomic UMAP (left) and DamID UMAP (right), colored by expression of pluripotency marker *Dppa5a*. **F**, Transcriptomic UMAP (left) and DamID UMAP (right), colored by expression of hematopoietic regulator *Tal1*. **G**, Genomic tracks of H3K27me3 and RING1B DamID signal per cluster at the *Tal1* locus. **H**, Heatmaps showing the H3K27me3 (left) and RING1B (right) DamID signal of all identified PRC targets for transcriptional clusters 3, 0, 1, 6, and 4. PRC targets are ordered based on hierarchical clustering. **I**, Fold-change in expression of Polycomb targets between clusters where the gene is PRC- associated and clusters where the gene is PRC-free.

We next sought to define the set of genes that is Polycomb-regulated in the EB system. First, we determined the H3K27me3 and RING1B signal at the promoter region of all genes (Methods) and compared these two readouts across the clusters. This confirmed good correspondence between H3K27me3 and RING1B profiles (Fig. S3F-G), albeit with a slightly higher signal amplitude for RING1B (Fig. S3G). This difference between RING1B and H3K27me3 may be biological (e.g., differential binding sites or kinetics) and/or technical (e.g., the use of a full-length protein versus a mintbody to target Dam). Nonetheless, because of the overall similarity between the two profiles, we decided to classify high-confidence Polycomb targets as having both H3K27me3 and RING1B enrichment in at least one of the EB clusters (excluding cluster 7 due to the relatively low number of cells) or in the previous ESC data set (Methods). We identified 9,159 Polycomb-regulated targets across the entire dataset, in good concordance with a previous study in mouse development (Gorkin et al., 2020) (Fig. S3H). These results validate the quality of the EpiDamID data and underscore the potential of the method to derive cell type-specific chromatin profiles from complex tissues.

Next, we intersected the cluster-specific transcriptome and DamID data to relate gene expression patterns to Polycomb associations. Based on the known role of Polycomb in gene silencing, differential binding of PRC1/2 to genes is expected to be associated with changes in expression levels of these genes. As exemplified in Fig. 3F-G, the cell type-specific expression of *Tal1*, a master regulator in hematopoiesis, in cluster 6 is indeed associated with an absence of H3K37me3 and RING1B exclusively in this cluster, whereas strong Polycomb occupancy over the *Tal1* promoter is evident in all other clusters. For presentation purposes, we display only the most prominent pluripotent cluster (3) and omit the other pluripotent clusters (2, 5, and 7), which had very similar characteristics. The negative association of Polycomb binding with gene expression is apparent for all PRC targets that are upregulated in the hematopoietic cluster (Fig. S3I-J). In addition, unsupervised clustering of H3K27me3 and RING1B promoter occupancy across cell clusters shows variation in signal between target genes as well as between cell types, indicating dynamic regulation of these targets within the EB system (Fig. 3H). Moreover, the subset of Polycomb targets that shows variable PRC occupancy is typically more highly expressed in the clusters where Polycomb is absent (Fig. 3I). Collectively, these data illustrate the strength of the EpiDamID approach to capture transcription and chromatin dynamics during differentiation in a single integrated method.

### Polycomb-regulated transcription factors form separate regulatory networks

After confirming the validity of the EpiDamID approach in measuring Polycomb dynamics during differentiation, we next focused on the Polycomb targets based on their function. We found that TF genes are over-represented within the Polycomb target genes (Fig. S4A), in line with previous observations (Boyer et al., 2006). Nearly half of all TF genes in the genome (761/1689) is bound by Polycomb in at least one cluster. In addition, genes encoding TFs generally accumulate higher levels of H3K27me3 and RING1B compared to other protein-coding genes (Fig. S4B). In line with an important role in lineage specification, Polycomb-controlled TFs are expressed in a cell type-specific pattern, as opposed to the more constitutive expression across cell types for Polycomb-independent TFs (Fig. S4C-D). Accordingly, the Polycomb-controlled TFs are enriched for Gene Ontology (GO) terms associated with animal development (Fig. S4E).

The enriched Polycomb targeting of developmentally regulated TF genes inspired us to further investigate the role of Polycomb in TF network hierarchies. We used SCENIC to systematically identify target genes that are associated with the expression of TFs (Aibar et al., 2017; van de Sande et al., 2020). SCENIC employs co-expression patterns as well as binding motifs to link TFs to their targets, together henceforth termed “regulons” (per SCENIC nomenclature). We identified 285 “activating” regulons after filtering (Fig. 4A and Methods). Notably, while regulons and their activity were found independently of RNA-based cluster annotations, we observed excellent recapitulation of cluster-specific transcriptional networks, confirming that SCENIC-identified regulon activity holds information on cellular identity (Fig. 4A).

**Figure 4.**
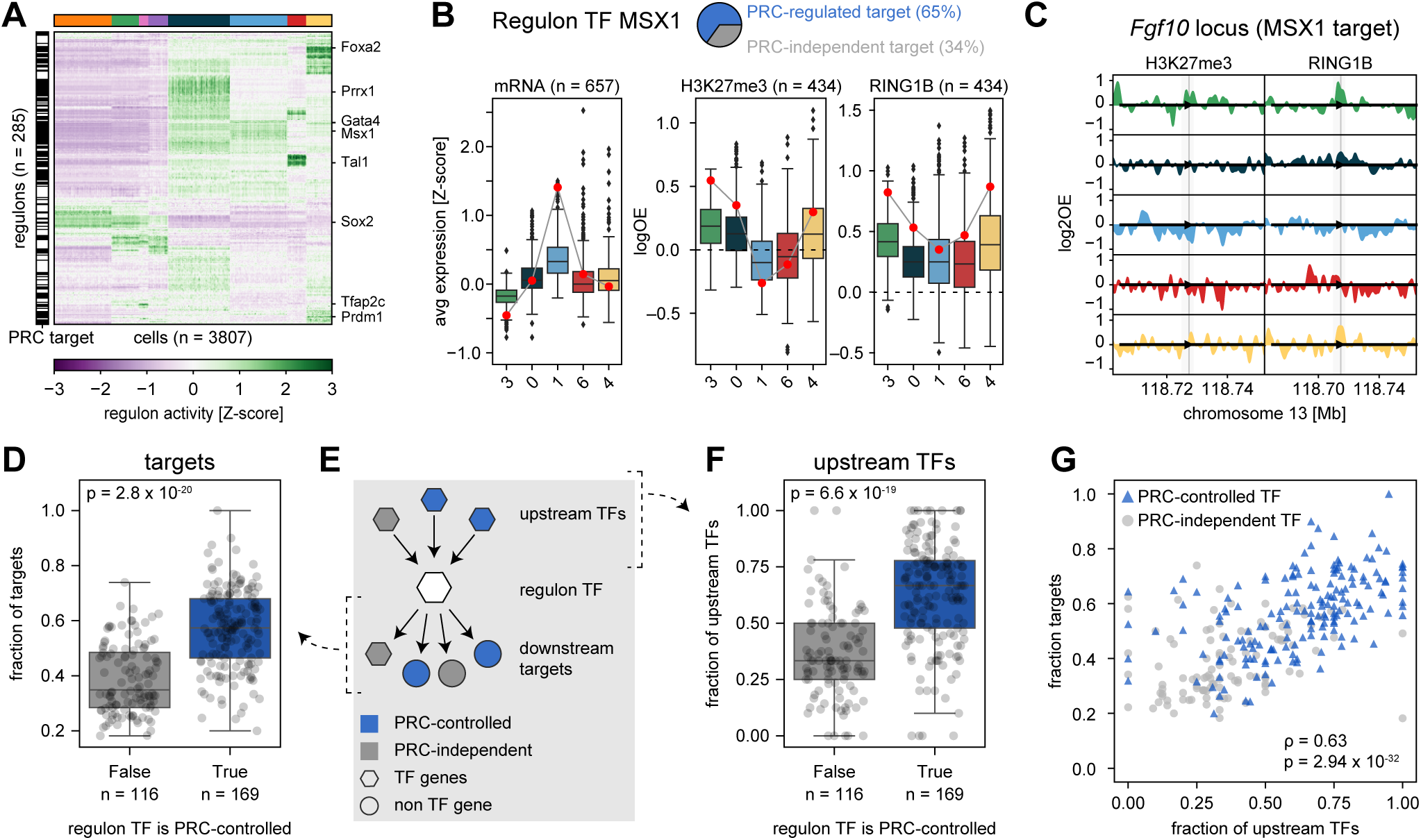
Polycomb-regulated transcription factors form separate regulatory networks. **A**, Heatmap showing SCENIC regulon activity per single cell. Cells (columns) are ordered by transcriptional cluster; regulon (rows) are ordered by hierarchical clustering. The black and white bar on the left indicates whether the regulon TF is a PRC target (black) or not (white). **B**, Example of the relationship between expression and Polycomb regulation for the MSX1 regulon. The pie chart indicates the percentages of Polycomb-controlled or Polycomb-independent target genes (blue and grey, respectively). Left: boxplots showing target gene expression (averaged *Z*-score) per cluster for all target genes. Middle and right: boxplots showing the H3K27me3 and RING1B DamID signal at the TSS per cluster for the Polycomb-controlled target genes. The expression and DamID signal of *Msx1* is indicated with a red circle. **C**, Genomic tracks of H3K27me3 and RING1B DamID signal per cluster at the *Fgf10* locus, one of the target genes of MSX1. **D**, Boxplots showing the fraction of Polycomb-controlled target genes, split by whether the TF itself is Polycomb-controlled. **E,** Schematic of the regulatory network, indicating the relationship between a regulon TF (white hexagon), its upstream regulators (colored hexagons), and its downstream targets (colored hexagons/circles). **F**, Boxplots showing the fraction of Polycomb-controlled upstream regulators, split by whether the regulon TF is Polycomb-controlled. **G,** Scatter plot showing the relationship between the fraction of Polycomb-controlled targets and regulators of a regulon TF. Regulon TFs that are PRC controlled are indicated in blue; regulon TFs that are PRC independent are indicated in grey. Correlation was computed using Spearman’s rank correlation.

Based on the expression modules identified with SCENIC, we first sought to determine how overall regulon activity correlates to Polycomb binding. As illustrated for the homeobox TF gene *Msx1*, we found that regulon activity is generally anti-correlated with Polycomb association of both the TF gene (red dot) and its Polycomb-controlled target genes (boxplots, 65% of all MSX1 targets) (Fig. 4B-C). We wondered whether there is a general preference for Polycomb-controlled TFs to target genes that themselves are regulated by Polycomb. This is indeed the case: while Polycomb-controlled TFs have a similar number of target genes compared to other TFs (Fig. S4F), the expression of the targets is much more frequently controlled by Polycomb than would be expected by chance (Fig. 4D). This effect is even stronger when considering the subset of targets that is exclusively regulated by Polycomb TFs (Fig. S4G). Using the transcriptional network provided by SCENIC, we also identified upstream TFs that control the expression of the regulon TFs (Fig. 4E). Similar to the target genes, the regulators of Polycomb-controlled TFs also tend to be Polycomb-controlled (Fig. 4F). Moreover, the fractions of Polycomb-controlled upstream regulators and downstream targets are correlated (Fig. 4G), indicating consistency in the level of Polycomb regulation across at least three layers of the TF network. Since Polycomb plays an important role in cell type specification, we evaluated whether this strict Polycomb control in the network was exclusive to lineage-specific genes. By clustering TFs into lineage-specific and unspecific groups based on their expression pattern (Fig. S4H), we found that, while this trend was especially strong for the lineage-specific genes, the consistency of Polycomb regulation in the network was a feature for other, unspecific, genes as well (Fig. S4I). These results suggest that Polycomb-associated hierarchies exist, forming relatively separate networks isolated from other gene regulatory mechanisms, and that this phenomenon extends beyond lineage-specific genes alone.

Together, the above findings demonstrate that single-cell EpiDamID can be successfully applied in complex developmental systems to gather detailed information on cell type-specific Polycomb regulation and its interaction with transcriptional networks.

### Implementation of EpiDamID during zebrafish embryogenesis

As presented above, DamID is a method that requires insertion of the Dam fusion protein into the biological system of interest. Genetic engineering of cell lines offers the advantage of many powerful *in vitro* differentiation systems, exemplified by our work in EBs. Contrastingly, it has proven challenging to apply DamID as a tool to study embryogenesis in transgenic vertebrate model organisms. To overcome this limitation, we previously established a protocol that introduces DamID into mouse preimplantation embryos via microinjections in the zygote (Borsos et al., 2019; Pal et al., 2021). Here, we sought to implement a similar strategy to apply EpiDamID during zebrafish development.

To establish the system, we profiled heterochromatin marked by H3K9me3 in single cells. H3K9me3 is reprogrammed during the early stages of development in several species (Laue et al., 2019; Mutlu et al., 2018; Rudolph et al., 2007; Santos et al., 2005; C. Wang et al., 2018) and the deposition of this mark coincides with decreased developmental potential (Ahmed et al., 2010). It was previously shown in zebrafish that H3K9me3 is largely absent before the maternal-to-zygotic transition (MZT) due to the presence of maternal *smarca2* mRNA. Upon zygotic transcription, degradation of *smarca2* results in a gradual increase of H3K9me3 from MZT up to shield stage [6 hours post-fertilization (hpf)] (Laue et al., 2019). However, it remains unclear whether the H3K9me3 distribution undergoes further remodeling after this stage, and whether its establishment differs across cell types during development.

To address these questions and to test EpiDamID in a zebrafish developmental context, we used the MPHOSPH8 chromodomain targeting H3K9me3 (Kungulovski et al., 2014), which we validated for EpiDamID in RPE-1 cells (Fig. 1B, F). We injected *Dam-Mphosph8* mRNA into the yolk at 1-cell stage (Fig. 5A) and optimized the mRNA concentrations to obtain scDam&T-seq data of high quality (data not shown). We separately injected *Dam-Mphosph8* to profile H3K9me3, and untethered *Dam* to profile chromatin accessibility. Embryos were collected at the 15-somite stage, which comprises a wide diversity of cell types corresponding to all germ layers. We generated 2,127 single-cell samples passing both DamID and CEL- Seq2 thresholds (Methods). To validate the specificity of the obtained H3K9me3 signal, we combined the DamID data of all cells in an *in silico* whole-embryo sample and compared this to the published H3K9me3 ChIP-seq data of 6-hpf embryos (Laue et al., 2019), which showed good concordance (Fig. S5.1A). These data confirm that we have established an orthogonal non-transgenic approach to generate high-resolution genome-wide profiles of heterochromatin during zebrafish development.

**Figure 5.**
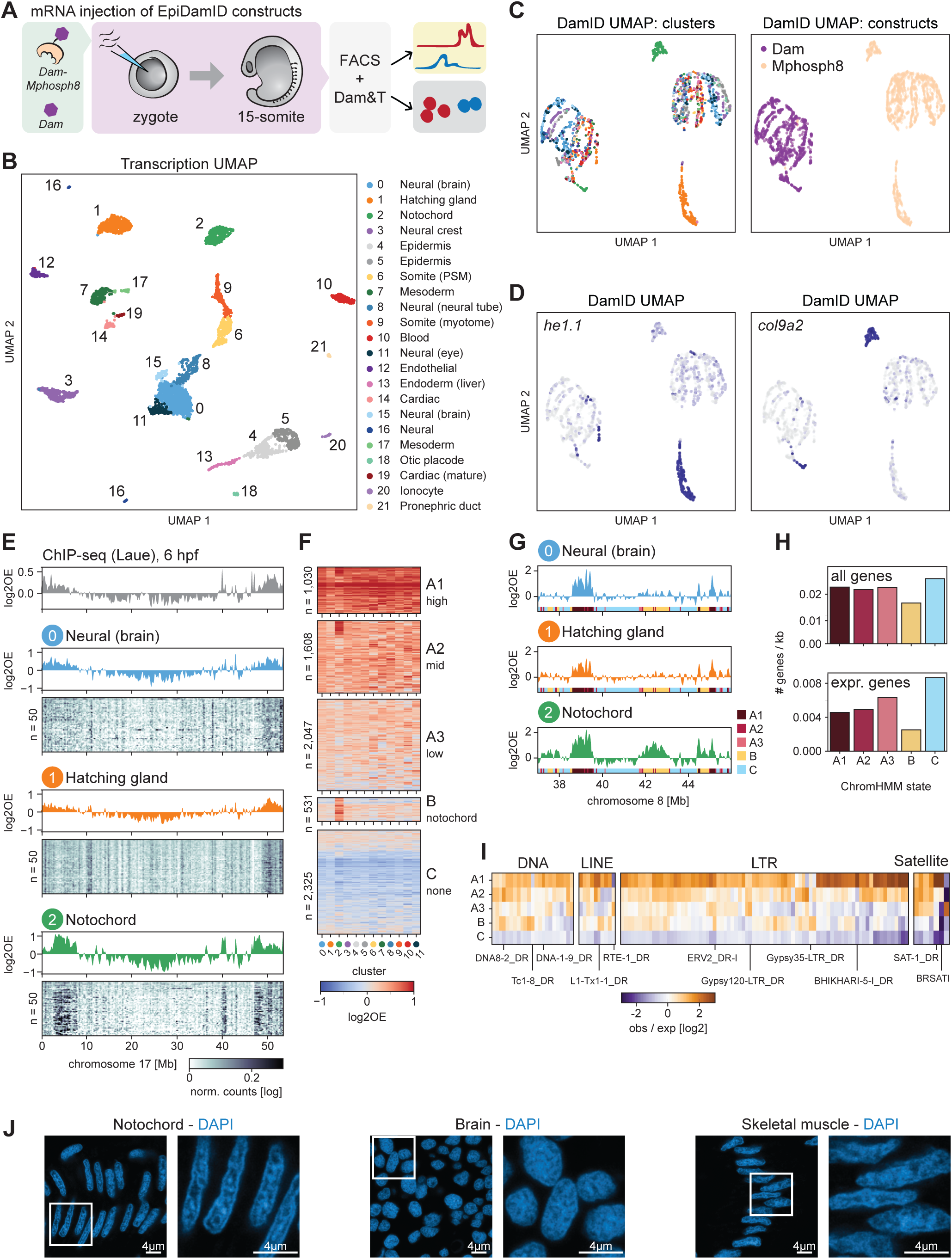
Notochord-specific H3K9me3 enrichment in the zebrafish embryo. **A**, Schematic representation of the experimental design and workflow. **B**, UMAP based on the transcriptional readout of all single-cell samples passing CEL-Seq2 thresholds (n = 3902). **C**, UMAP based on the genomic readout of all single-cell samples passing DamID thresholds (n = 2833). Samples are colored by transcriptional cluster (left) and Dam-targeting domain (right). **D**, Expression of the hatching gland marker *he1.1* (left) and the notochord marker *col9a2* (right) projected onto the DamID UMAP. **E**, Genomic H3K9me3 signal over chromosome 17. Top track: H3K9me3 ChIP-seq signal normalized for input control from the 6-hpf embryo, taken from Click or tap here to enter text.. Remaining tracks: Combined single-cell DamID MPHOSPH8 data normalized for the Dam control for clusters 0-2. **F**, Heatmap showing the cluster-specific average H3K9me3 enrichment over all domains called per ChromHMM state. Only clusters with >30 single-cell MPHOSPH8 and Dam samples were used for the ChromHMM (i.e., clusters 0-11). Per state, the domains have been clustered using hierarchical clustering. **G**, Genomic H3K9me3 signal over a part of chromosome 8 for clusters 0-2. The colored regions at the bottom of each track indicate the ChromHMM state. **H**, Gene density of all genes (top) and expressed genes (bottom) per state. **I**, Enrichment of repeats over all different ChromHMM states. Only repeats having at least 100 copies throughout the genome and an enrichment ≥2 in at least one state are shown. Enrichment is computed as the observed number of repeats in a state compared to the expected number based on the genome coverage of that state. Example repeats are indicated. **J,** Representative confocal microscopy images of DAPI staining in cryosections of notochord (left), brain (middle), and skeletal muscle (right) of 15-somite zebrafish embryos.

### Broad domains of notochord-specific H3K9me3 enrichment revealed by scDam&T-seq

Analysis of the single-cell transcriptome data resulted in 22 clusters of diverse cell types (Fig. 5B), which we annotated according to expression of known marker genes (Fig. S5.1B). We performed dimensionality reduction based on the DamID signal and observed a clear separation of cells according to their Dam construct, and to a lesser extent on their cell type (Fig. 5C-D). As described for EpiDamID in EBs, cluster-specific DamID profiles allowed us to employ the LDA classifier to recover a further 705 cells (Fig. S5.2C). Notably, the MPHOSPH8 samples of hatching gland (cluster 1, *he1.1* expression) and notochord (cluster 2, *col9a2* expression) segregated strongly from the other cell types (Fig. 5D), implying differences in their single-cell H3K9me3 profiles. Subsequently, we combined all single-cell DamID data per cluster to obtain cell type-specific H3K9me3 patterns (Fig. 5E). We observed clear differences in the genomic profiles, most notably the appearance of large domains of H3K9me3 enrichment in the notochord, and overall lower levels of H3K9me3 in the hatching gland (Fig. 5E and Fig. S5.1D). These differences are largely consistent between individual cells (Fig. 5E, heatmaps). In conclusion, with EpiDamID we are able to capture cell type-specific epigenetic profiles for individual cells of the 15-somite zebrafish embryo.

Next, to more systematically identify and characterize regions of differential H3K9me3 enrichment between cell clusters, we performed ChromHMM (Ernst & Kellis, 2012, 2017) (Methods). The approach uses the H3K9me3 signal per cluster to annotate genomic segments as belonging to different H3K9me3 “states” characterized by the clusters in which the segment is enriched. We included the 12 cell clusters with sufficient cells (containing >30 cells per construct) and identified five H3K9me3 states across the genome. These represented: A) three states of constitutive H3K9me3 with different enrichment levels [A1-A3], B) notochord-specific H3K9me3 enrichment, and C) constitutive depletion of H3K9me3 (Fig. 5F-G). State A (A1-3) chromatin forms broad domains (Fig. S5.1E) that together comprise 27% of the genome (Fig. S5.1F) and, as expected for H3K9me3-associated chromatin regions, are characterized by sparser gene density and lower gene activity compared to the H3K9me3- depleted state C (Fig. 5H). Moreover, state A1 is strongly enriched for zinc-finger transcription factors (Fig. S5.1G), which are known to be demarcated by H3K9me3 in other species (Hahn et al., 2011). The notochord-specific state B has similar characteristics to states A1-A3 (Fig. 5H, S5.1E-F), yet exhibits broader consecutive regions of H3K9me3 enrichment (Fig. 5G and S5.1E) and an even lower active gene density (Fig. 5H). Despite the size of the notochord-specific H3K9me3 domains and their features typical of repressive chromatin, we could not relate them back to differences in tissue-specific gene expression (Fig. S5.1H).

One of the known functions of H3K9me3 chromatin is the repression of transposable elements (Bulut-Karslioglu et al., 2014; S. Liu et al., 2014; Mosch et al., 2011). Indeed, it was previously observed in zebrafish that nearly all H3K9me3 domains in early embryos are associated with repeats (Laue et al., 2019). We therefore determined whether distinct repeat classes were over-represented in each H3K9me3 ChromHMM state (Fig. S5.2A). This analysis revealed a strong enrichment of several repeat classes in state A1, including LTR and tRNA. To further discriminate within the classes, we looked into all repeat types with at least 100 genomic copies. In addition to the most prominent enrichment of LTR repeats, state A1 showed a high frequency of pericentromeric satellite repeats SAT-1 and BRSATI (Fig. 5I), in line with the known occupancy of H3K9me3 at pericentromeric regions in diverse species. We postulated that the enrichment of repeats found within state A1, as opposed to A2 and A3, could be due to elevated signal at these loci, as a result of active targeting for H3K9me3- mediated silencing. Inspection of the DamID patterns indeed showed a clear increase of signal centered on specific repeat regions in state A1, and to lesser extents in other states (Fig. S5.2B). In addition, we investigated whether there are repeats strongly enriched within the notochord-specific state B domains that could potentially explain the existence of these domains. We found that there are indeed certain repeats specifically enriched within state B (Fig. 5I and Fig. S5.2C), albeit rarely as conspicuous as the enrichments observed in state A1. It therefore warrants further study to see whether H3K9me3 is involved in cell type-specific repression of repetitive genomic regions in notochord.

### Altered expression of chromatin proteins and pronounced nuclear compartmentalization in notochord

Finally, we took advantage of the combined measurements of transcriptome and epigenetic profiles to gain insight into cluster-specific expression of known chromatin proteins in relation to the differential H3K9me3 patterns. We inspected the expression of histone methyltransferases, demethylases and other chromatin factors across clusters, and did not detect an upregulation of known H3K9 methyltransferases (*setdb2*, *setdb1a/b*, *suv39h11a/b*, *ehmt2*) nor demethylases (*kdm4aa/ab/b/c*, *phf8*) in notochord (Fig. S5.2D). Of note, however, the H3K9- and H3K36-specific demethylase *kdm4c* was exclusively upregulated in hatching gland, which could explain the low H3K9me3 levels in this cluster. Moreover, among genes significantly upregulated in notochord, *lmna* stood out. This gene encodes the nuclear lamina protein Lamin A/C that is known to associate with heterochromatin (Gruenbaum & Foisner, 2015) and plays an important structural role in the nucleus (Donnaloja et al., 2020; Gruenbaum & Foisner, 2015), further suggesting an altered chromatin state in the notochord.

To directly investigate chromatin state and nuclear organization in these embryos, we performed confocal imaging of H3K9me3 and DAPI stainings in notochord, brain, and skeletal muscle. H3K9me3-marked chromatin displayed a typical nuclear distribution in all tissues, including heterochromatin foci as previously reported (Laue et al., 2019) (Fig. S5.2E). Notably, the DAPI staining showed markedly more structure in the notochord compared to the other tissues (Fig. 5J), visible as a clear rim along the nuclear periphery and denser foci within the nuclear interior. Since DAPI is indicative of AT-rich and generally less accessible DNA, this suggests a stronger separation between euchromatin and heterochromatin. Together with our findings of notochord-specific H3K9me3 domains and differential expression of chromatin factors, these observations on nuclear organization warrant further study of their contribution to the structural properties of notochord cells.

Collectively, the implementation of EpiDamID in zebrafish embryos shows that this strategy provides a flexible and accessible approach to generate rich single-cell information on the epigenetic states that underlie biological processes during zebrafish embryogenesis.

## Discussion

Here, we have developed and tested EpiDamID, an adaptation of conventional DamID that extends its application of profiling DNA-protein contacts to epigenetic marks. EpiDamID utilizes the binding specificities of genetically encoded mintbodies (Sato et al., 2013, 2016; Tjalsma et al., 2021), histone PTM-identifying domains (Kungulovski et al., 2014, 2016; Vermeulen et al., 2007), or full-length chromatin readers to target Dam to specific chromatin marks. We presented a wide diversity of histone PTM patterns generated after viral transduction of EpiDamID constructs in RPE-1 cells, and validated the approach through comparison to genomic profiles generated with ChIP-seq (Fig. 1). A selection of histone PTMs was chosen to illustrate that EpiDamID yields high-quality single-cell profiles in engineered mESC lines (Fig. 2), which was further demonstrated by its implementation in an EB differentiation system (Fig. 3 and 4). Lastly, we showed that single-cell histone PTM profiling can be achieved in zebrafish embryos (Fig. 5). Joint single-cell quantifications of histone PTMs and transcriptome enabled the identification of cell types and associated histone PTM profiles in integrated experiments.

### Advantages of DamID for studying histone PTMs during embryogenesis

Since DamID was first developed (Vogel et al., 2007), a wide range of derivative technologies have been established (see (Aughey et al., 2019)). This includes the possibility to perform live-cell imaging of DamID-marked chromatin regions (Altemose et al., 2020; Borsos et al., 2019; Kind et al., 2013), targeted DamID (TaDa) for tissue-specific profiling without cell isolation or dissection (Marshall & Brand, 2017; Southall et al., 2013), proteomics on DamID-marked genomic regions (Wong et al., 2021), single-cell experiments (Altemose et al., 2020; Borsos et al., 2019; Kind et al., 2015; Rooijers et al., 2019), and protocols for performing DamID in living mouse preimplantation embryos (Borsos et al., 2019; Pal et al., 2021). Moreover, DamID has been successfully established in many model systems including various mouse and human cell lines and several organisms, including plants, fish, fly and worms (Aughey et al., 2019). EpiDamID can be implemented in any DamID-based protocol, thereby offering the possibility to obtain live-cell microscopic, proteomic and genomic measurements of histone PTMs in a single integrated toolbox in diverse biological settings. (Kind et al., 2013; Park et al., 2019).

The variety of implementations and model systems makes EpiDamID especially suitable to study histone PTMs in development. The single-cell implementations of DamID—scDamID and scDam&T-seq—require little sample handling and few enzymatic steps, resulting in minimal sample loss. This makes them particularly efficient and, as a result, offers the possibility to individually collect and process all cells belonging to the same tissue (Borsos et al., 2019). For example, scDam&T-seq with EpiDamID constructs could be used to individually collect all cells of a single preimplantation mouse embryo and examine epigenetic and transcriptomic differences that may point towards cell fate commitment. In contrast, state-of-the-art methods require extensive tissue handling prior to signal amplification, preventing capture and tracking of intra-embryonic variability. Furthermore, the DamID genomic marks are stable upon deposition, offering the interesting possibility to track ancestral EpiDamID genomic signatures through mitosis to study inheritance and spatial distribution of epigenetic states in daughter cells (Kind et al., 2013; Park et al., 2019). This feature adds a temporal axis to genomic experiments, albeit only for a single cell division due to the dilution of the ^m6^A-print upon DNA replication. This unique aspect of DamID warrants further exploration, especially in single-cell multimodal omics experiments.

### DamID as an integrative method for single-cell multi-modal omics

The DamID workflow is suitable for integration with other single-cell protocols to achieve multi-modal measurements (Markodimitraki et al., 2020). The limited handling prior to individual cell capture offers opportunities to integrate upstream steps of other protocols that are compatible with the final processing steps of scDamID. Powerful future method integrations may involve combining scDamID with scChIC-seq (Ku et al., 2019) or sortChIC (Zeller et al., 2021) to measure two genome-wide profiles in the same cell, or the incorporation of the CITE-seq approach to obtain quantitative single-cell measurements of protein abundance, transcriptomics and histone PTM profiles. This latter combination offers the exciting prospect to study gene expression across the central dogma of gene regulation.

### Chromatin-associated gene regulatory networks in mouse developmental systems

We implemented EpiDamID in EB cultures to investigate the PRC2-deposited H3K27me3 mark alongside the binding of core PRC1 component RING1B, and the role of Polycomb in transcriptional regulation during differentiation. We observed extensive variability in Polycomb occupancy across distinct cell types, and, in addition, identified the existence of hierarchical Polycomb-associated regulatory networks. We speculate that these Polycomb-controlled networks ensure robustness of stable maintenance of repression. By establishing the EpiDamID approach, we have set the stage for similar experiments in more dynamic biological systems, such as gastruloid cultures or *in vivo* mouse experiments. EpiDamID can be performed via mRNA injection to study early development as we have demonstrated here and previously (Borsos et al., 2019; Pal et al., 2021), or via the establishment of transgenic animals that conditionally express EpiDamID constructs.

### Structural function of notochord during zebrafish embryogenesis may be supported by heterochromatin organization in the nucleus

Lastly, we established a protocol to apply EpiDamID in zebrafish embryos. We identified broad regions of notochord-specific H3K9me3 enrichment with no evident function in cell-type specific gene silencing. The notochord is an important embryonic structure that serves a mechanical as well as a cellular signalling function to its surrounding tissues (Corallo et al., 2015). During embryogenesis, notochord cells develop a large vacuole, in which high osmotic pressure forces the tissue into its characteristic stack-of-coins appearance, and provides unique mechanical properties essential for the elongation of the embryo. We hypothesize that notochord-specific domains of H3K9me3 enrichment identified in our study may contribute to the unique structural properties of these cells, potentially to withstand the strong osmotic forces acting upon the nucleus. Since H3K9me3 is known to convey nuclear stiffness Click or tap here to enter text. broad domains of consecutive heterochromatin would be beneficial. Additional support for this possibility is provided by our observation of consistently increased mRNA levels of the *lmna* gene, encoding Lamin A/C, in notochord cells. Lamin A/C is a constituent of the nuclear lamina (NL), a filamentous network lining the inner nuclear membrane (Gruenbaum & Foisner, 2015) that is directly connected to the cytoskeleton via transmembrane proteins in the LINC complex (Crisp et al., 2006). Of note, the A-type Lamins are specifically responsible for modulating nuclear structure and rigidity (Donnaloja et al., 2020). Finally, the existence of an altered chromatin state in notochord is further supported by the observation that DAPI-stained DNA forms more clearly segregated structures in notochord cells compared to other cell types, implying a stronger separation between euchromatin and heterochromatin. These findings warrant further investigation into the nature of notochord heterochromatin and its role in supporting the structural properties of this tissue.

### Limitations

The limitations of EpiDamID are similar to those of DamID in general. In order to generate histone PTM profiles with EpiDamID, a construct encoding for the Dam-fusion protein needs to be expressed in the system of interest. This may involve a substantial time investment depending on the system of choice. Then, to achieve optimal signal over noise, the conditions to be optimized generally differ dependent on the properties of the Dam-fusion protein. For cell lines, it typically involves establishing a conditional expression system and performing a number of experiments to test for optimal induction times. For microinjection experiments, it requires optimizing mRNA concentrations that are injected in the zygote. The optimal mRNA concentration to achieve best signal-to-noise ratio depends on the histone PTM of interest and the developmental stage of choice. Finally, due to the *in vivo* expression and consequent roaming of the Dam-POI in the nucleus, spurious methylation gradually accumulates in unspecific, mostly accessible, chromatin regions. The degree of accumulated background signal differs substantially between different Dam-POIs, yet interferes most with proteins that reside within active chromatin. This can be overcome either computationally through normalization to the untethered Dam protein or the implementation of Dam mutants with decreased affinity for DNA. Unfortunately, we found that the reduced enzymatic activity of these mutants (Dam126 and others, data not shown) results in insufficient ^m6^A-events for high-quality single-cell profiling. Further adaptation of the Dam protein to achieve an enzyme with combined full enzymatic activity and reduced DNA-binding affinity may further improve the quality of EpiDamID profiles in single cells.

We expect that the study of chromatin associations in the context of dynamic cellular states will provide better understanding of the occurrence and identity of the events that shape chromatin-regulated gene expression, as well as functions outside of transcription.

## Materials availability

Unique material generated in this study is available from the Lead Contact with a completed Materials Transfer Agreement.

## Data and code availability

All data generated in this manuscript is deposited on the NCBI Gene Expression Omnibus (GEO) portal under accession number GSE184032. All custom code is available from the corresponding author (JK) upon request.

## Acknowledgements

We would like to thank the members of the Kind laboratory for their critical reading of the manuscript and helpful comments and suggestions. In particular, we thank Koos Rooijers for providing input and support on the computational work. This work was funded by an ERC Starting grant (ERC-StG 678423-EpiID) to JK. The Oncode Institute is partially funded by the KWF Dutch Cancer Society. IG is supported by an EMBO Long-Term Fellowship ALTF1214-2016, Swiss National Science Fund grant P400PB_186758 and NWO-ZonMW Veni grant VI.Veni.202.073. PDN is supported by an EMBO Long-Term Fellowship ALTF1129-2015, HFSPO Fellowship (LT001404/2017-L) and an NWO-ZonMW Veni grant (016.186.017-3). The laboratory of JB is supported by the Netherlands Cardiovascular Research Initiative: An initiative with support of the Dutch Heart Foundation and Hartekind, CVON2019-002 OUTREACH. The laboratory of HK is supported by MEXT/JSPS KAKENHI (JP18H05527, JP20K06484, and JP21H04764), and Japan Science and Technology Agency (JPMJCR16G1 and JPMJCR20S6). We additionally thank the Hubrecht Sorting Facility, the Hubrecht Imaging Center, and the Utrecht Sequencing Facility (USEQ) subsidized by the University Medical Center Utrecht.

## Author contributions

FJR, KLdL, SSdV and JK conceived and designed the study. FJR, KLdL and JK wrote the manuscript. FJR performed all computational analyses. KLdL and SSdV performed all experiments unless noted otherwise. CVQ and EB designed and generated knock-in mouse ESC lines. PDN performed all zebrafish experiments, with assistance from IG and SSdV, and supervised by JB. YS and HK generated materials. All authors discussed the results and contributed to the final manuscript.

## Declaration of interest

The authors declare no competing interests.

## Figure legends

**Figure S1.**
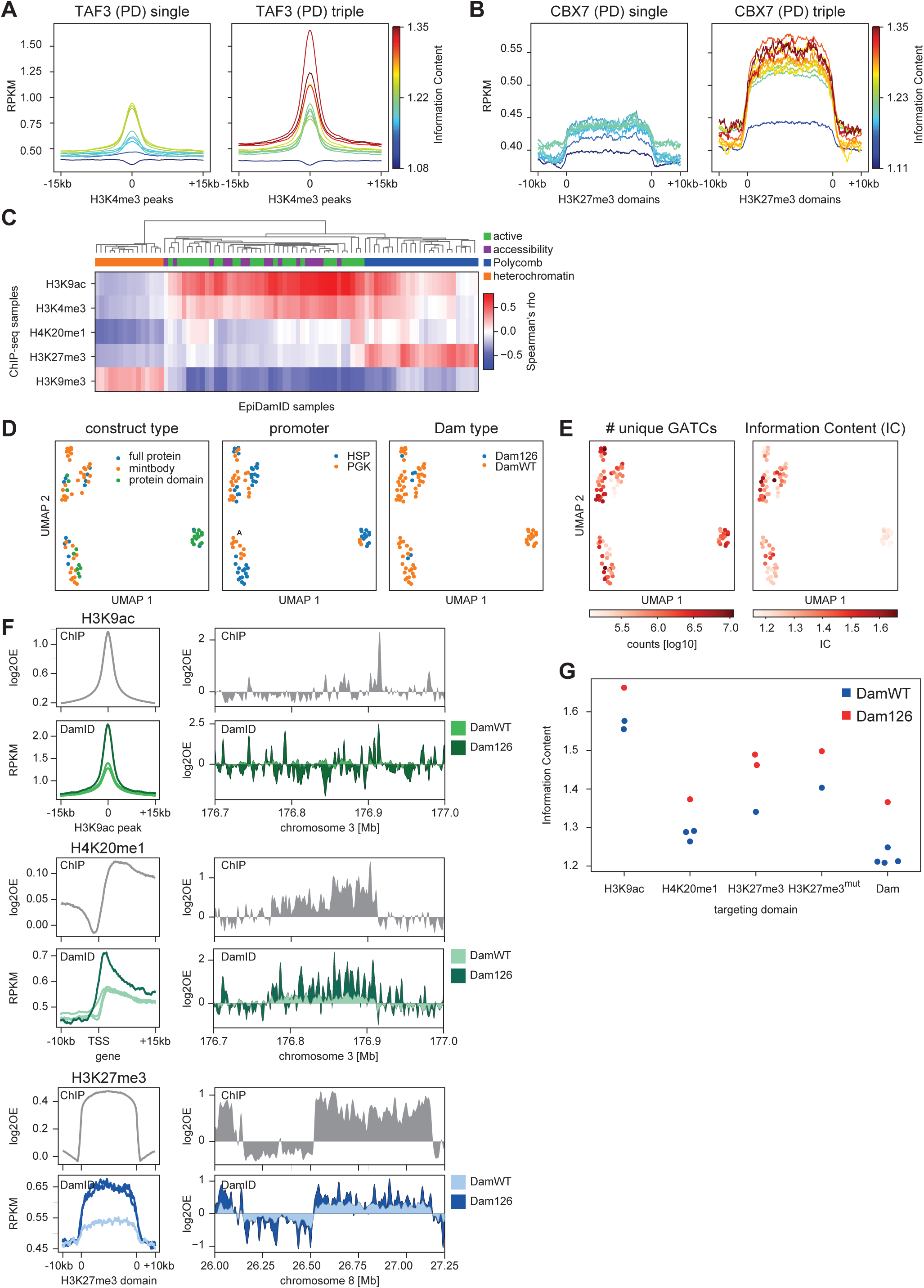
Technical validation of EpiDamID data. **A-B**, Average enrichment over genomic regions of interest for TAF3 (**A**) and CBX7 (**B**) DamID. Left: data generated by fusing Dam to a single protein domain; Right: data generated by fusing Dam to a trimer of the same protein domain. Sample lines are colored by their Information Content (IC). **C**, Clustered heatmap showing the correlation between DamID and ChIP-seq samples. Correlations were computed using Spearman’s rank correlation. **D**, UMAPs of samples, colored by construct properties. **E**, UMAPs of samples, colored by DamID-seq depth and Information Content. **F,** Left, average DamID and ChIP-seq enrichment plots over genomic regions of interest. Signal is normalized for untethered Dam and input, respectively. Regions are the TSS of genes for H3K9ac (top), gene bodies for H4K20me1 (middle), and ChIP-seq domains for H3K27me3 (bottom). **F**, Right, genome browser view of ChIP-seq and DamID enrichment corresponding to left panels. The data shown in D-F represents the combined data of all samples of a particular targeting domain.

**Figure S2:**
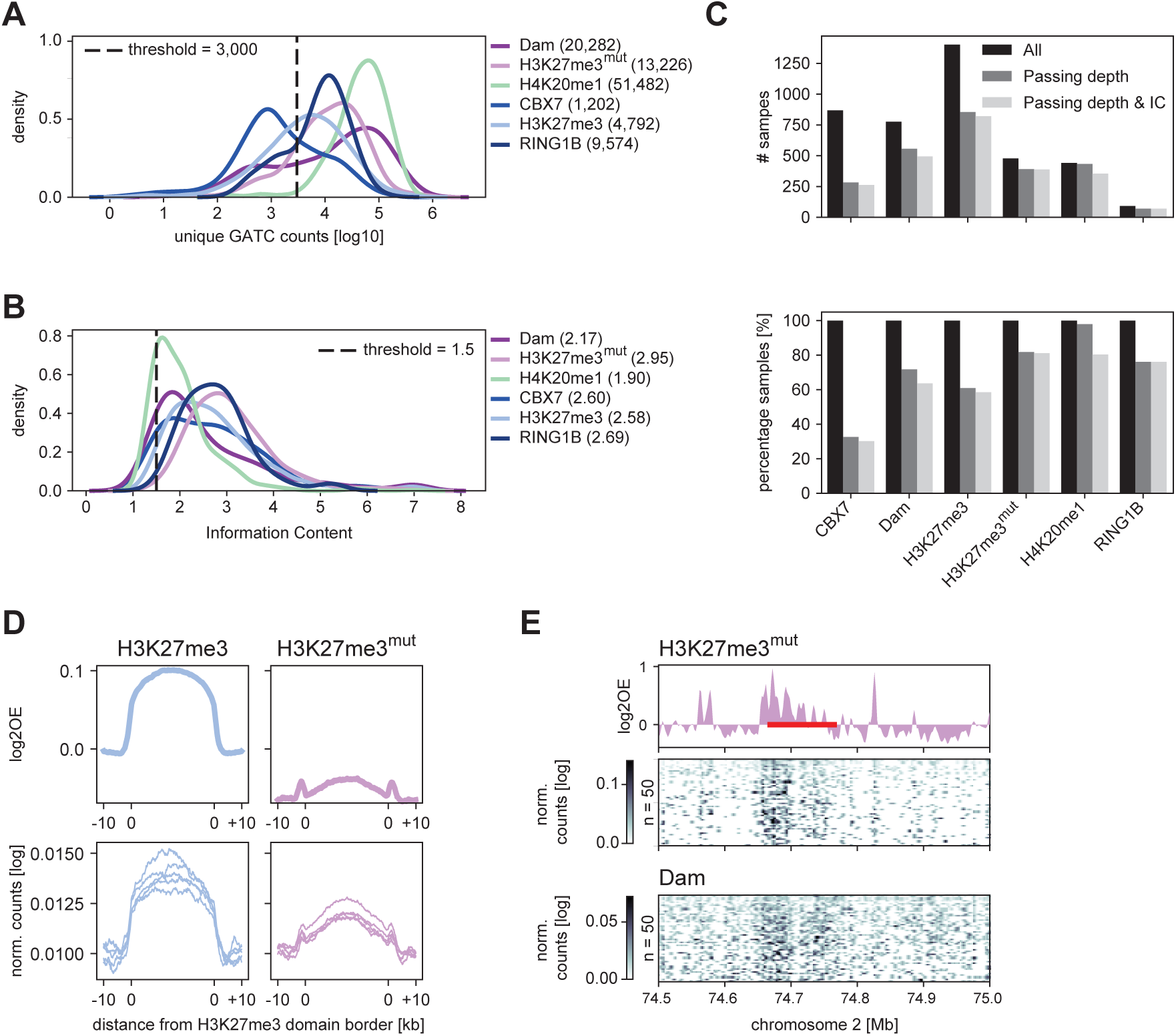
Detection of histone PTMs in single mouse embryonic stem cells with a single-cell implementation of EpiDamID. **A**, Density plot indicating the distribution of the number of unique GATCs detected for each cell line. The dashed line indicates the threshold used for data filtering. **B**, Density plot indicating the distribution of the Information Content (IC) after filtering on depth for each cell line. The dashed line indicates the threshold used for data filtering. **C**, Overview of the number (top) and percentage (bottom) of samples retained after filtering on depth and IC. **D**, Average signal over H3K27me3 ChIP-seq domains of H3K27me3 and H3K27me3^mut^ mintbodies. Top: *in silico* populations normalized for Dam; Bottom: five of the best single-cell samples (bottom) normalized by read depth. **E**, Signal of H3K27me3^mut^ and Dam control over the *HoxD* cluster and neighboring regions. The DamID track show the Dam-normalized *in silico* populations of H3K27me3^mut^, while the heatmaps show the depth-normalized single-cell data of the fifty richest cells for H3K27me3^mut^ and Dam. The red bar around 74.7 Mb indicates the *HoxD* cluster.

**Figure S3.**
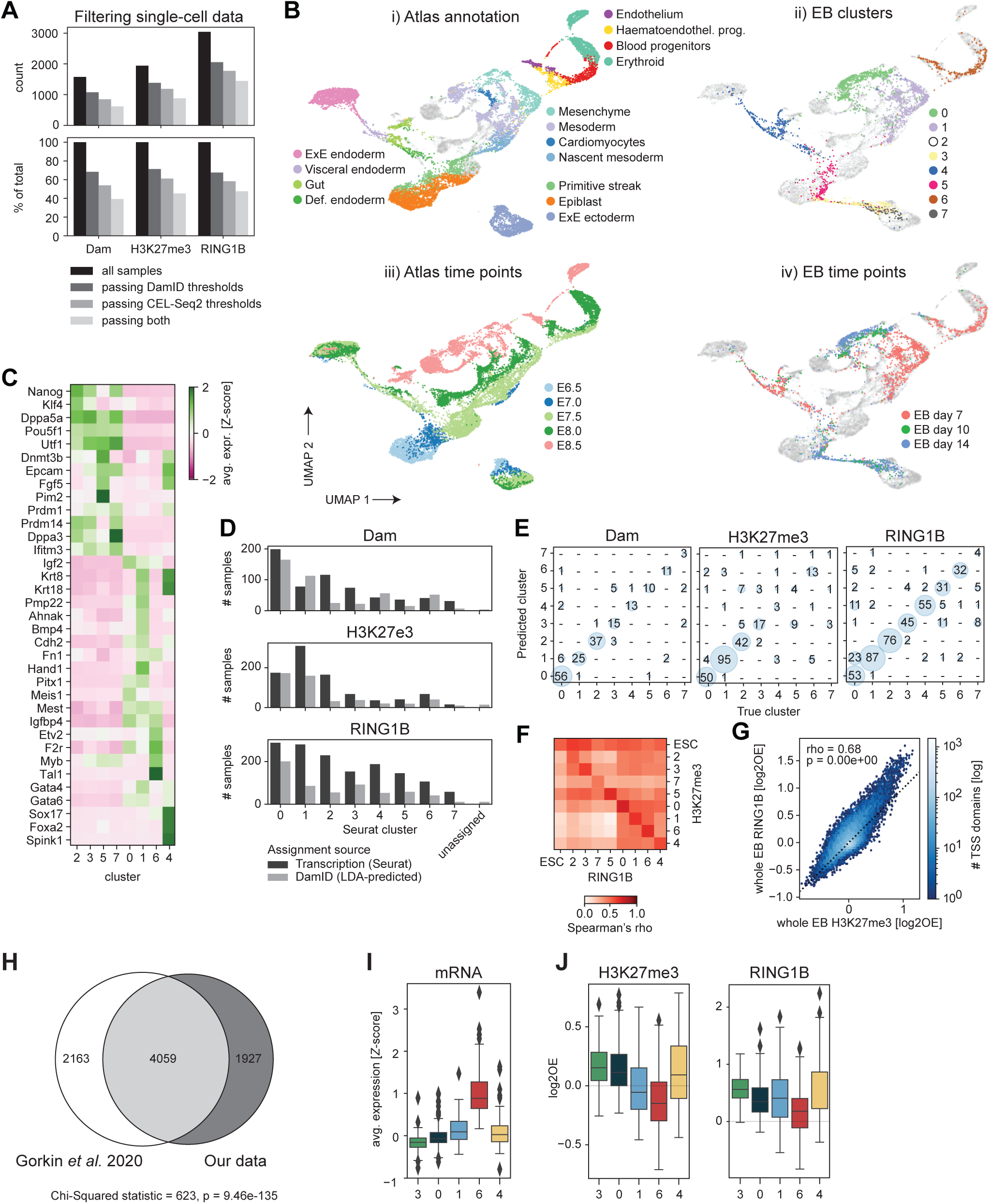
Validation and characterization of scDam&T-seq data in mouse embryoid bodies. **A,** Overview of the number (top) and percentage (bottom) of remaining samples after application of DamID and/or CEL-Seq2 filtering. **B,** UMAPs of samples based on the integration of our EB transcription data with single-cell RNA-seq mouse embryonic data Click or tap here to enter text., colored by reference-annotated cell type (i), EB-annotated cluster (ii), atlas embryonic day (iii) and EB day (iv). For atlas integration, the day 0 (i.e., mESC) time point was excluded. **C,** Average expression of known marker genes. Expression was standardized over single-cells and the per-cluster average was computed. **D,** Bar plots showing the number of cells per cluster assigned by Seurat (i.e. based on transcriptional readout) or the LDA classifier (i.e. based on DamID readout). **E,** Confusion plots showing the performance of the LDA classifier during training, for each construct. **F,** Correlation between the combined H3K27me3 and RING1B DamID signal at the TSS of all genes per transcriptional cluster. **G,** Correlation of combined H3K27me3 and RING1B DamID signal at the TSS of all genes. Data of all single-cell samples passing DamID thresholds was combined for each construct. **H,** Overlap between a published set of PRC targets during mouse development Click or tap here to enter text. and our PRC targets. Significance of the overlap was computed with a Chi-squared test. **I**, Boxplots showing the expression (averaged *Z*-score) of genes identified as significantly upregulated in cluster 6. **J,** Boxplots showing the H3K27me3 (left) and RING1B (right) DamID signal at the TSS of the subset of genes shown in **I** that are PRC targets.

**Figure S4.**
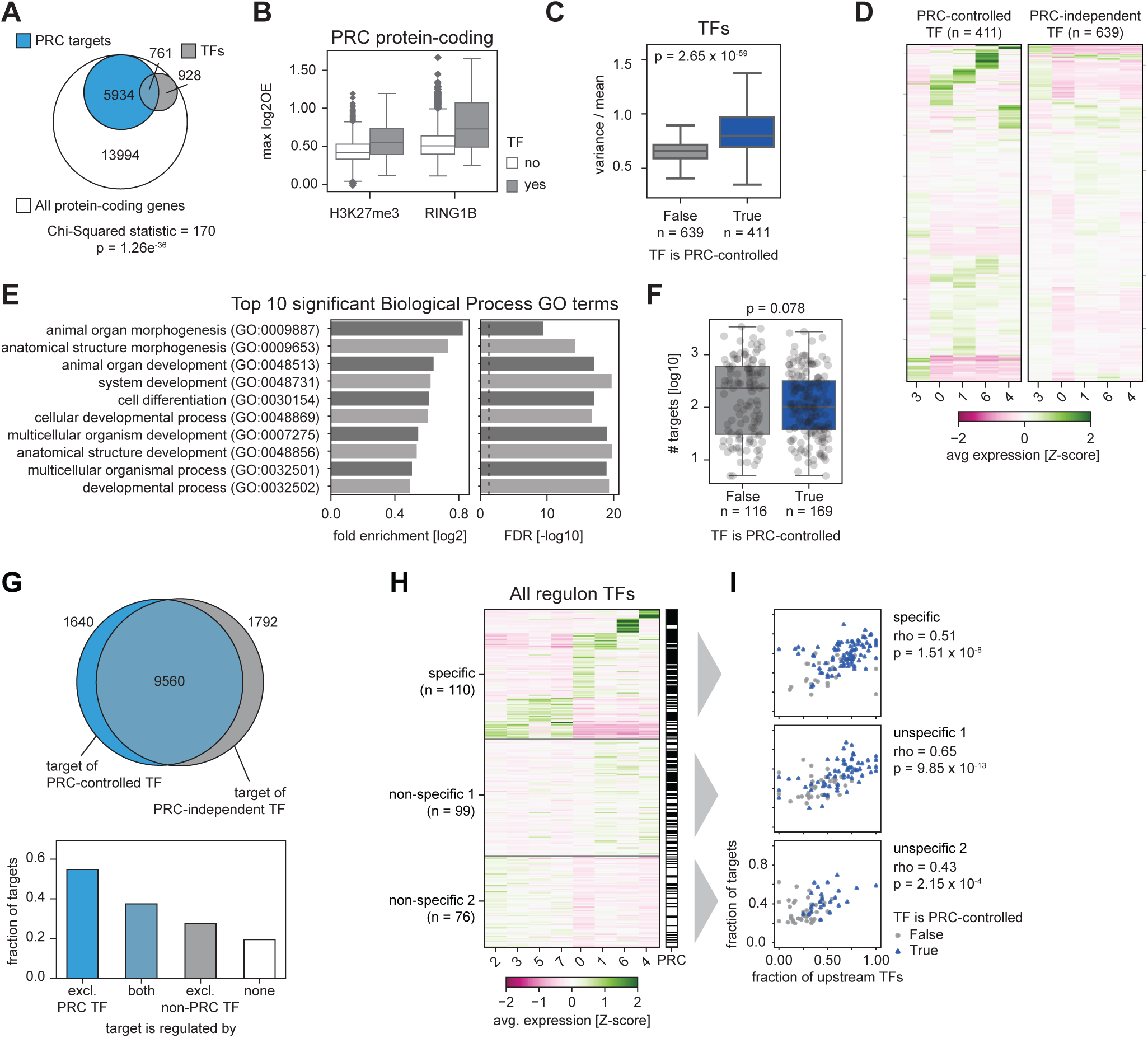
Characterization of the Polycomb-regulated regulatory network. **A,** Venn diagram showing the overlap between PRC-controlled protein-coding genes (blue) and transcription factors (TF) (grey) in the context of all protein-coding genes (white). The significance of the overlap between PRC targets and TFs was computed using a Chi-squared test. **B,** Boxplots showing the maximum observed H3K27me3 and RING1B DamID signal across transcriptional clusters for PRC-controlled TFs (grey) and the remaining PRC-controlled protein-coding genes (white). **C**, Quantification of variability in gene expression of PRC-regulated and PRC-independent TFs (only expressed genes are included). Boxplots show variance over mean across all single cells. Significance was computed using a Mann-Whitney-U test. **D**, Clustered heatmaps showing mRNA expression (averaged Z-score) per cluster, of Polycomb-regulated TFs (left) and Polycomb-independent TFs (right). Only expressed genes are included in this plot. **E,** The ten most significant Biological Process GO terms between PRC-controlled and PRC-independent TFs. **F**, Number of targets of each regulon TF, split by whether or not the TF is PRC-regulated. Difference between categories was tested with a Mann-Whitney-U test. **G,** Top: Venn diagram displaying the overlap between genes that are targets of a PRC- controlled TF (blue) and genes that are targets of a PRC-independent TF (grey). **G,** Bottom: Bar plot showing the fraction of targets in each category that is PRC-regulated. **H,** Clustered heatmap showing mRNA expression (averaged *Z-*score) per cluster, of all regulon TFs, grouped by lineage-specific or non-specific genes. TFs are annotated as PRC- controlled (black) or PRC independent (white). **I**, Scatter plot showing the relationship between the fraction of Polycomb-controlled targets and regulators of a regulon TF. Regulon TFs that are PRC controlled are indicated in blue; regulon TFs that are PRC independent are indicated in grey. Regulon TFs are split based on the groups indicated in **H**. Correlation was computed using Spearman’s rank correlation.

**Figure S5.1:**
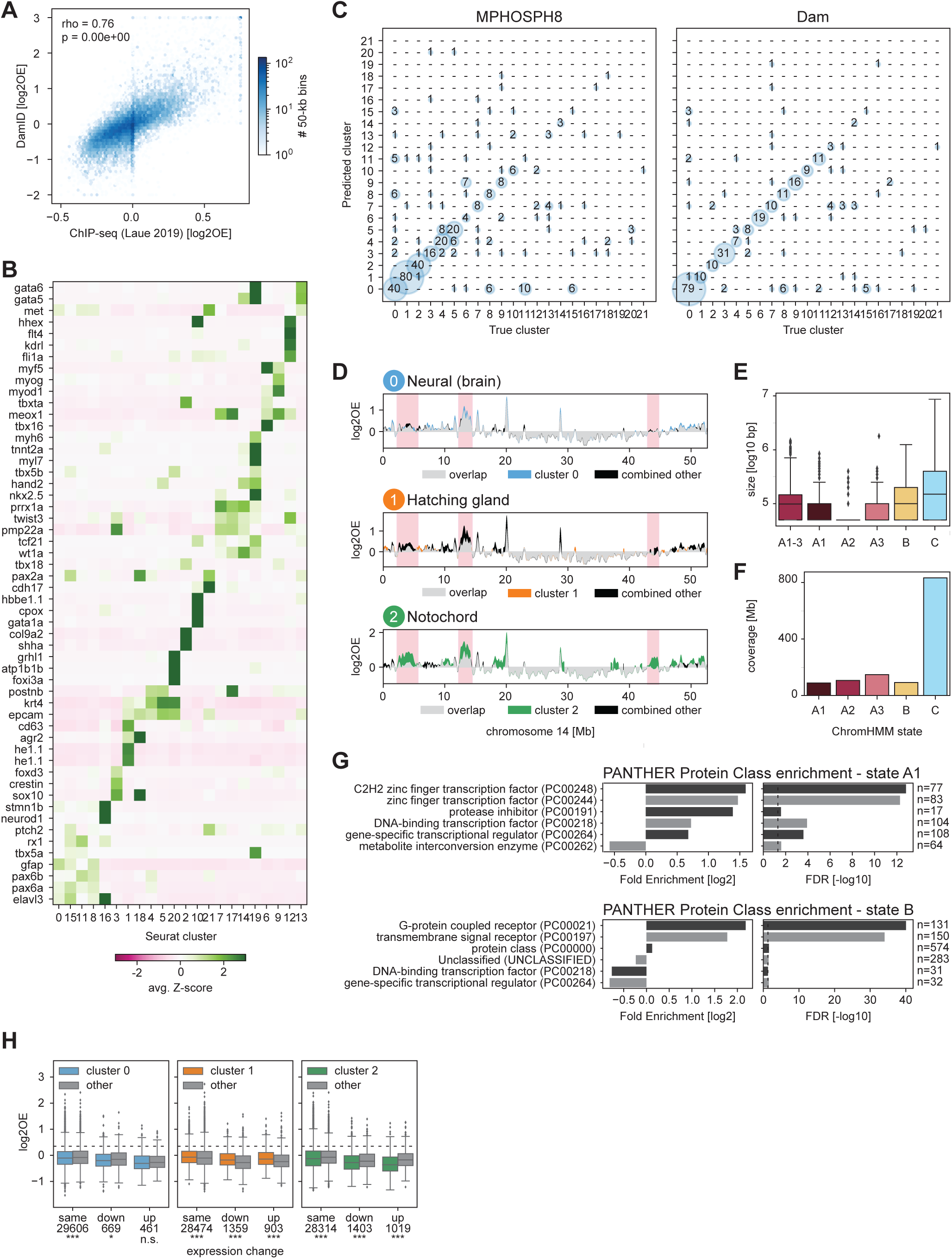
Characterization of transcriptomic clusters and associated genomic H3K9me3 enrichments. **A**, Comparison of our data with a published H3K9me3 ChIP-seq dataset of the 6-hpf zebrafish embryo Click or tap here to enter text.. All single-cell MPHOSPH8 and Dam samples were combined to generate an *in silico* whole-embryo data set; DamID data is the log2OE of MPHOSPH8 signal over Dam is shown; ChIP-seq is the log2OE of H3K9me3 over input control. **B**, Expression of marker genes over all clusters, ordered by cell type. The average single-cell Z-scores are shown. **C,** Confusion plots showing the performance of the LDA classifier during training, for each construct. **D,** Genomic H3K9me3 signal over chromosome 14. For clusters 0-2, the cluster-specific signal (color) is compared to the combined signal from all other clusters (black). Each set indicates the overlay, where overlapping regions are colored grey. **E**, Distribution of domain sizes per ChromHMM state and for states A1-3 combined. **F,** Total genomic coverage per ChromHMM state. **G,** PANTHER protein-class enrichments Click or tap here to enter text. for genes found in state A1 (top) and B (bottom). **H,** H3K9me3 enrichment at differentially expressed genes for each cluster. Each boxplot shows for upregulated, downregulated and stable genes the H3K9me3 signal of the corresponding cluster and the combined signal of the complementary clusters. The significance of the difference in H3K9me3 was tested with a Mann-Whitney-U test, *** indicates a p-values smaller than 0.001, ** a p-value smaller than 0.01, * a p-value smaller than 0.1, and n.s. a p-value larger than 0.1.

**Figure S5.2:**
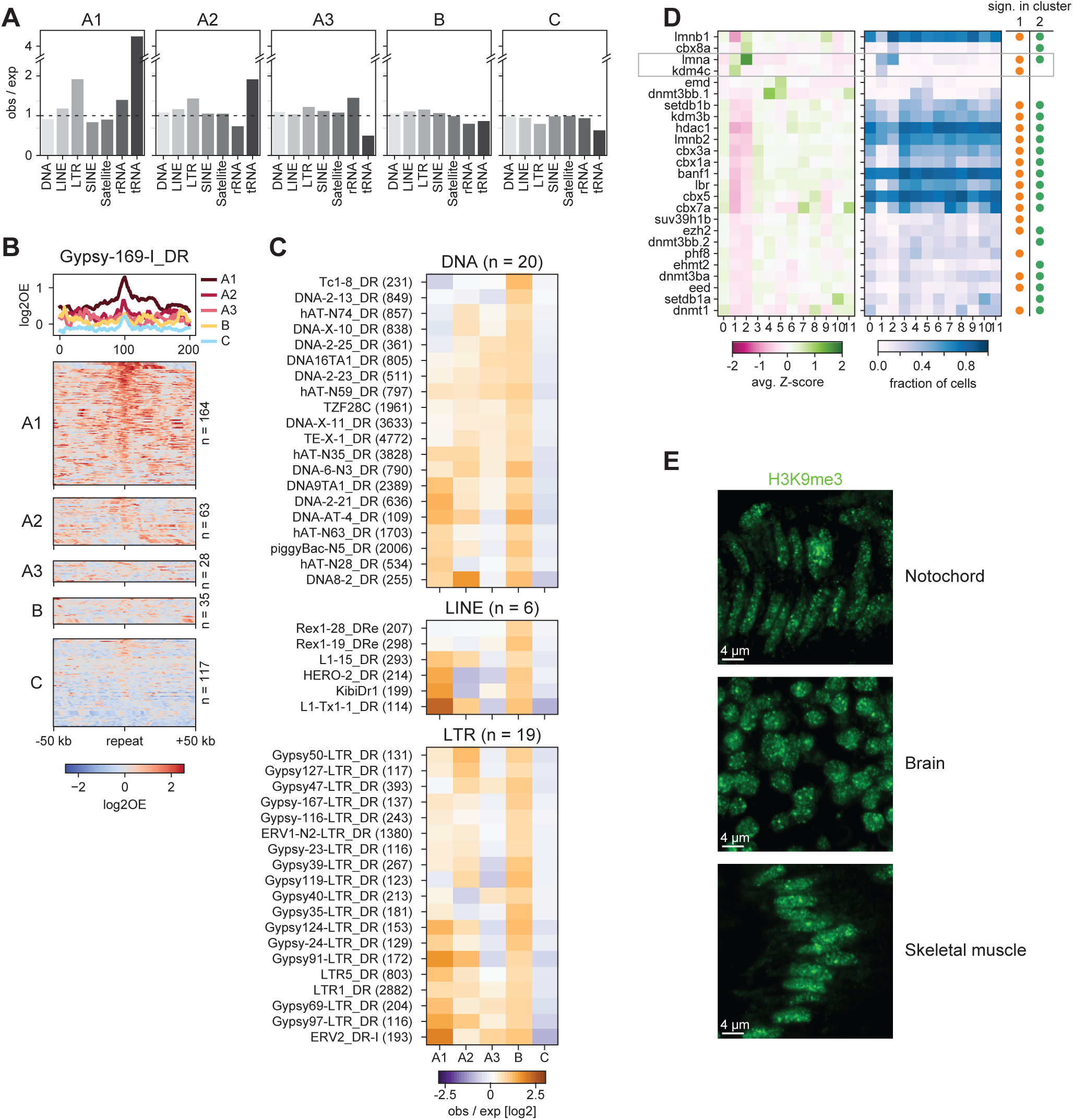
Characterization of repeat content, expression of chromatin factors and nuclear localization of H3K9me3 chromatin. **A**, Enrichment of repeats per class for all ChromHMM states. Enrichment is computed as the observed number of repeats within a state relative to the expected number based on the genome coverage of each state. **B**, H3K9me3 enrichment at Gypsy-169-I_DR repeats across ChromHMM states. The heatmaps show the enrichment per individual repeat instance, while the line plot shows the average enrichment per state. **C**, Enrichment of repeats in ChromHMM states as in Figure 5I. Only repeats having at least 100 copies throughout the genome and an enrichment ≥1.5 in state B are included. Enrichment is computed as the observed number of repeats in a stated compared to the expected number based on the genome coverage of that state. **D**, Expression of various chromatin factors across clusters 0-11. The left heatmap shows the average single-cell expression (Z-score); the right heatmaps shows the fraction of cells in each cluster with at least one transcript of each gene. Only factors that are expressed in at least 10% of cells of at least one cluster are shown. **F**, Representative images of H3K9me3 staining in cryo-sections of notochord (left), brain (middle), and skeletal muscle (right) in 15-somite embryos.

## Methods

### Chromatin immunoprecipation followed by high-throughput sequencing

Mouse embryonic stem cells and hTERT-immortalized RPE-1 cells were cultured following ATCC instructions. ChIP-seq was performed as described previously (Collas, 2011), with the following adaptations. Cells were harvested by trypsinization, and chemically crosslinked with fresh formaldehyde solution (1% in PBS) for 8 minutes while rotating at room temperature. Crosslinking was quenched with glycine on ice and sample was centrifuged at 500 g for 10 min at 4 °C. Pellet was then resuspended in lysis buffer for 5 min on ice and sonicated as follows: 16 cycles of 30 s on / 30 s off at max power (Bioruptor Diagenode), and centrifuged at 14,000 rpm at 4 °C for 10 min. The chromatin in supernatant was treated with RNase A for 30 min at 37 °C, and Proteinase K for 4 hours at 65 °C to reverse crosslinks, then cleared using DNA purification columns and eluted in nuclease-free water. Chromatin was incubated with antibodies (see below), after which Protein G beads (ThermoFisher #88847) were added for antibody binding. After successive washing, samples were cleared using DNA purification columns, eluted in nuclease-free water, and measured using a Qubit fluorometer. Libraries were prepared according to the Illumina TruSeq DNA LT kit and sequenced on the Illumina HiSeq 2500 following manufacturer’s protocols. Up to 50 ng of immunoprecipitated chromatin was used as input for library preparation. Antibodies used were: anti-H3K4me3 Abcam ab8580, anti-H3K9ac Abcam ab4441, anti-H3K9me3 Abcam ab8898, anti-H3K27me3 Merck Millipore 07-449, anti-H3K36me3 Active Motif 61902, anti-H4K20me1 Abcam ab9051.

### Lentiviral DamID construct design and production

The constructs for mintbodies, chromatin binding domains, and full-length protein constructs were fused to Dam in both possible orientations under the control of the auxin-inducible degron (AID) system (Kubota et al., 2013; Nishimura et al., 2009) with either the hPGK or HSP promoter, and cloned into the pCCL.sin.cPPT.ΔLNGFR.Wpre lentiviral construct (Amendola et al., 2005) by standard cloning procedures. Lentivirus was produced as previously described (Amendola et al., 2005) and the PGK viruses were concentrated approximately 40-fold using Amicon Ultra-15 centrifugal filter units (Merck #UFC910024), the HSP expressed viruses were used unconcentrated.

### Bulk DamID2

hTERT-RPE1 cells were grown in DMEM/F12 (Gibco) containing 10% FBS (Sigma F7524 lot BCBW6329) and 1% Pen/Strep (Gibco) in 6-well plates. At 30% confluence, cells were transduced with 1500 μL total volume unconcentrated lentivirus, amounts ranging between 20-1500 μL unconcentrated lentivirus (or 0.1-40 μL concentrated) in the presence of 10 μg/mL polybrene. Cells were collected for genomic DNA isolation (Wizard, Promega) 48 h after transduction. Dam methylation levels were checked by ^m6^A-PCR as previously described (de Luca & Kind, 2021; Vogel et al., 2007) and sequenced following the DamID2 protocol (Markodimitraki et al., 2020).

### Immunofluorescent staining and confocal imaging

Viral transduction was performed as described above for bulk DamID2, with the exception that RPE-1 cells were grown on glass coverslips. Two days after transduction, cells were washed with PBS and chemically crosslinked with fresh formaldehyde solution (2% in PBS) for 10 minutes at RT, permeabilized (with 0.5% IGEPAL® CA-630 in PBS) for 20 minutes and blocked (with 1% bovine serum albumin (BSA) in PBS) for 30 minutes. All antibody incubations were performed in final 1% BSA in PBS followed by three PBS washes at RT. Incubation with primary antibody against the endogenous histone modification as well as purified ^m6^A-Tracer protein (Schaik et al., 2020) (recognizing methylated DNA) was performed at 4 °C for 16 hours (overnight), followed by anti-GFP (against ^m6^A-Tracer protein) incubation at RT for 1 hour, and secondary antibody incubations at RT for 1 hour. The final PBS wash was simultaneously an incubation with DAPI at 0.5 μg/mL for 2 min, followed by a wash in MilliQ and sample mounting on glass slides using VECTASHIELD Antifade mounting medium (Vector Laboratories). Primary antibodies: anti-H3K9ac abcam ab4441 (rabbit) at 1:1000, anti-H3K9me3 abcam ab8898 (rabbit) at 1:300, anti-GFP Aves GFP-1020 (chicken) at 1:1000. Secondary antibodies: AlexaFluor anti-chicken 488 at 1:500 and anti-rabbit 647 at 1:500. Imaging was performed on a Leica SP8 confocal microscope with a 63x (NA 1.40) oil-immersion objective. Images were processed in Imaris 9.3 (Bitplane) by baseline subtraction. Additional background correction was done with a 1-μM Gaussian filter for the images of Dam-CBX1 ^m6^A-Tracer and H3K9me3 stainings.

### Generation of mouse embryonic stem cell lines

The various stable clonal F1 hybrid mESC lines for the initial single cell experiments were created by lentiviral co-transduction of pCCL-EF1α-Tir1-IRES-puro and pCCL-hPGK-AID-Dam-POI constructs with a 4:1 ratio in a EF1α-Tir1-IRES-neo mother line (Rooijers et al., 2019), after which the cells were selected for 10 days on 0.1% gelatine coated 10-cm dishes in 60% Buffalo Rat Liver (BRL)-conditioned medium containing 0.8 μg/mL puromycin (Sigma P9620), 250 μg/mL G418 (ThermoFisher 11811031) and 0.5 mM IAA. Individual puromycin resistant colonies were handpicked and tested for the presence of the constructs by PCR using Dam specific primers fw-ttcaacaaaagccaggatcc and rev-gacagcggtgcataaggcgg.

The clonal F1 hybrid knock-in cell lines were CRISPR targeted in a mother line carrying Tir1-Puro in the TIGRE locus (Zeng et al., 2008). For all CRISPR targeting, cells were cultured on gelatin-coated 6-wells in 60% BRL conditioned medium to 70-90% confluency and transfected with Lipofectamin3000 (Invitrogen L3000008) according to the supplier protocol with 2 μg donor vector and 1 μg Cas9/guide vector. At 24 h after transfection the cells were split to a gelatin-coated 10-cm dish and antibiotic selection of transfected cells is started 48 h after transfection. Cells were selected with 60% BRL conditioned medium containing 0.8 µg/mL puromycin for the Tir1 knock-in and 2.5 µg/mL blasticidin (Invivogen) for the AID-Dam knock-in lines. After 5-10 days of selection, individual colonies were manually picked and screened by PCR for the correct genotype.

All CRISPR knockin lines were made in a Tir1-TIGR mother line that was generated by co-transfection of Cas9-gRNA plasmid pX330-EN1201(Addgene plasmid #92144) and donor plasmid pEN396-pCAGGS-Tir1-V5-2A-PuroR TIGRE (Addgene plasmid #92142) (Nora et al., 2017). The Tir1-puro clones were screened for the presence of Tir1 by PCR from the CAGG promoter to Tir1 with the primers fw-cctctgctaaccatgttcatg and rev-tccttcacagctgatcagcacc, followed by screening for correct integration in the TIGRE locus by PCR from the polyA to the TIGRE locus with primers fw-gggaagagaatagcaggcatgct and rev-accagccacttcaaagtggtacc. The Tir1 expression is further confirmed by Western blot using a V5 antibody (Invitrogen R960-25).

A knock-in of AID-Dam in the N-terminus of the RINGB1B locus was made by co-transfection of a donor vector carrying the blasticidin-p2A-HA-mAID-Dam cassette flanked by 2 500bp homology arms of the endogenous RING1B locus (pHom-BSD-p2A-HA-mAID-Dam) and p225a-RING1B spCas9-gRNA vector (sgRNA: 5’gctttttattcctagaaatgtctc3’) as described above. Picked clones were screened for correct integration by PCR with primers from Dam to the RING1B locus outside the targeting construct; fw-gaacaacaagcgcatctggc and rev-tcctcccctaacctgcttttgg. Presence of the RING1B wildtype allele was checked by PCR with primers fw-tcctcccctaacctgcttttgg and rev-gccttgcctgcttggtttg. The H3K27me3 mintbody coupled to ER-mAID-Dam was knocked into the Rosa26 locus by co-transfection of pHom- ER-mAID-V5-Dam-scFv_H3K27me3-P2A-BSD-Hom donor vector and p225a-Rosa26 spCas9-RNA vector (sgRNA: gtccagtctttctagaagatgggc) as described above. Picked clones were screened for correct integration by PCR from a sequence adjacent to the Rosa homology arm to the Rosa26 locus with primers fw-gaactccatatatgggctatg and rev-cttggtgcgtttgcgggga. The untethered mAID-Dam was knocked into the Rosa26 locus by co-transfection with the pHom-ER-mAID-V5-Dam-P2A-BSD-Hom donor vector and p225a-Rosa26 spCas9-RNA vector (sgRNA: gtccagtctttctagaagatgggc) as described above. Picked clones were screened for correct integration by PCR with the same primers as for the Dam-H3K27me3 mintbody knock-in line.

All clones with correct integrations were furthermore screened for their level of induction upon IAA removal by ^m6^A-PCR evaluated by gel electrophoresis (de Luca & Kind, 2021; Vogel et al., 2007), followed by DamID2 sequencing in bulk (Markodimitraki et al., 2020), to select the clone with a correct karyotype and the best signal to noise ratio of enrichment over expected regions or chromatin domains. Finally, the best 3-4 clones were selected for testing of IAA removal timing in single cells by DamID2.

### Mouse embryonic stem cell culture and induction of Dam-fusion proteins

F1 hybrid 129/Sv:Cast/Eij mouse embryonic stem cells (mESCs) were cultured on irradiated primary mouse embryonic fibroblasts (MEFs), in mESC culture media CM+/+ defined as follows: G-MEM (Gibco) supplemented with 10% FBS (Sigma F7524 lot BCBW6329), 1% Pen/Strep (Gibco), 1x GlutaMAX (Gibco), 1x non-essential amino acids (Gibco), 1x sodium pyruvate (Gibco), 0.1 mM β-mercaptoethanol (Sigma) and 1000 U/mL ESGROmLIF (EMD Millipore ESG1107). Cells were split every 3 days and medium was changed every other day. Expression of the Dam-POI constructs was suppressed by addition of 0.5 mM indole-3-acetic acid (IAA; Sigma, I5148). Lines were tested routinely for mycoplasma.

When plated for targeting or genomics experiments, cells were passaged at least 2 times in feeder-free conditions, on plates coated with 0.1% gelatin, grown in 60% BRL-conditioned medium, defined as follows and containing 1 mM IAA: 40% CM+/+ medium and 60% of CM+/+ medium conditioned on BRL cells. For timed induction of the constructs the IAA was washed out at different clone specific optimized times before single cell sorting.

### Embryoid body differentiation and induction of Dam-fusion proteins

For EB differentiation, the stable knock-in F1ES lines were cultured for 2 weeks on plates coated with 0.1% gelatin, grown in 2i+LIF ES cell culture medium defined as follows: 48% DMEM/F12 (Gibco) and 48% Neurobasal medium (Gibco), supplemented with 1x N2 (Gibco), 1x B27 supplement + vitamin A (Gibco), 1x non-essential amino acids, 1% FBS, 1% Pen/Strep, 0.1mM β-mercaptoethanol, 1 μM PD0325901 (Axon Medchem, PZ0162-5MG), 3 μM CHIR99021 (Tocris, SML1046-5MG), 1000 U/mL ESGRO mLIF. EB differentiation was performed according to ATCC protocol. On day 1 of differentiation, 2x10^6 cells were grown in suspension on a non-coated bacterial 10-cm dish with 15 mL CM +/- (with β-mercaptoethanol, without LIF) and 0.5 mM IAA. On day 2, half the cell suspension was divided over five non-coated bacterial 10-cm dishes each containing 15mL CM+/- medium and 0.5 mM IAA. Plates were refreshed every other day. EBs were harvested at day 7, 10, and 14. Two days before single-cell sorting, the EBs were grown in CM+/- medium containing 1 mM IAA, and induced as follows: 6 h without IAA (RING1B); 20 h without IAA and 7 h with 1 μM 4OHT (Sigma SML1666) (Dam-H3K27me3-mintbody); 7 h without IAA and 4 h with 1 μM 4OHT (untethered Dam). The EBs were evaluated by brightfield microscopy and hand-picked for further handling (see below).

### FACS for single-cell experiments

FACS was performed on BD FACSJazz or BD FACSInflux Cell Sorter systems with BD Sortware. mESCs and EBs were harvested by trypsinization, centrifuged at 300 g, resuspended in medium containing 20 μg/mL Hoechst 34580 (Sigma 63493) per 1x10^6^ cells and incubated for 45 minutes at 37°C. Prior to sorting, cells were passed through a 40-μm cell strainer. Propidium iodide (1 μg/mL) was used as a live/dead discriminant. Single cells were gated on forward and side scatters and Hoechst cell cycle profiles. Index information was recorded for all sorts. One cell per well was sorted into 384-well hard-shell plates (Biorad, HSP3801) containing 5 μL of filtered mineral oil (Sigma #69794) and 50 nL of 0.5 μM barcoded CEL-Seq2 primer (Markodimitraki et al., 2020; Rooijers et al., 2019). In the EB experiment, the knock-in mESC lines were cultured alongside on 2i+LIF medium and included as a reference at each timepoint.

### Single-cell Dam&T-seq

The scDam&T-seq protocol was performed as previously described in detail (Markodimitraki et al., 2020), with the adaptation that all volumes were halved to reduce costs. Liquid reagent dispension steps were performed on a Nanodrop II robot (Innovadyne Technologies / BioNex). Addition of barcoded adapters was done with a mosquito LV (SPT Labtech). In short, after FACS, 50 nL per well of lysis mix (0.07% IGEPAL, 1 mM dNTPs, 1:50,000 ERCC RNA spike-in mix (Ambion, 4456740)) was added, followed by incubation at 65 °C for 5 min. 100 nL of reverse transcription mix (1× First Strand Buffer and 10 mM DTT (Invitrogen, 18064-014), 2 U RNaseOUT Recombinant Ribonuclease Inhibitor (Invitrogen, 10777019), 10 U SuperscriptII (Invitrogen, 18064-014)) was added, followed by incubation at 42 °C for 2 h, 4 °C for 5 min and 70 °C for 10 min. Next, 885 nL of second strand synthesis mix (1× second strand buffer (Invitrogen, 10812014), 192 μM dNTPs, 0.006 U *E. coli* DNA ligase (Invitrogen, 18052019), 0.013 U RNase H (Invitrogen, 18021071), 0.26 U *E. coli* DNA polymerase (Invitrogen)) was added, followed by incubation at 16 °C for 2 h. 250 nL of protease mix was added (1× NEB CutSmart buffer, 1.0 mg/mL Proteinase K (Roche, 000000003115836001)), followed by incubation at 50 °C for 10 h and 80 °C for 20 min. Next, 115 nL of DpnI mix (1× NEB CutSmart buffer, 0.1 U NEB DpnI) was added, followed by incubation at 37 °C for 6 h and 80 °C for 20 min. Finally, 50 nL of 0.5uM DamID2 adapters were dispensed (final concentrations 25 nM), followed by 400 nL of ligation mix (1× T4 Ligase buffer (Roche, 10799009001), 0.13U T4 Ligase (Roche, 10799009001)) and incubation at 16°C for 16 hrand 65°C for 10 min. Contents of all wells were pooled and the aqueous phase was recovered by centrifugation and transfer to clean tubes. Samples were purified by incubation for 10 min with 0.8 volumes magnetic beads (CleanNA, CPCR-0050) diluted 1:7 with bead binding buffer (20% PEG8000, 2.5 M NaCl), washed twice with 80% ethanol and resuspended in 8 μl of nuclease-free water before in vitro transcription at 37 °C for 14 h using the MEGAScript T7 kit (Invitrogen, AM1334). Library preparation was done as described in the CEL-Seq2 protocol with minor adjustments (Hashimshony et al., 2016). Amplified RNA (aRNA) was purified with 0.8 volumes beads as described above, and resuspended in 20 μL of nuclease-free water, and fragmented at 94 °C for 90 sec with the addition of 0.25 volumes fragmentation buffer. Fragmentation was stopped by addition of 0.1 volumes of 0.5 M EDTA pH 8 and quenched on ice. Fragmented aRNA was purified with beads as described above, and resuspended in 12 μl of nuclease-free water. Thereafter, library preparation was done as previously described (Hashimshony et al., 2016) using up to 7 μL or approximately 150 ng of aRNA, and 8-10 PCR cycles depending on input material. Libraries were sequenced on the Illumina NextSeq500 (75-bp reads) or NextSeq2000 (100-bp reads) platform.

### Zebrafish

All animal experiments were conducted under the guidelines of the animal welfare committee of the Royal Netherlands Academy of Arts and Sciences (KNAW). Adult zebrafish (*Danio rerio*) were maintained and embryos raised and staged as previously described (Aleström et al., 2019; WESTERFIELD & M., 2000).

### Collection of zebrafish samples and FACS

Tübingen longfin (wild type) pairs were set up and the following morning, approximately 1 n L of 1 ng/μL *Dam-Mphosph8* mRNA or 0.5 ng/μL *Dam-Gfp* mRNA was injected into the yolk at the 1 cell stage. Embryos were slowed down overnight at 23°C and the following morning all embryos were manually dechorionated. At 15-somite stage, embryos were transferred to 2- mL Eppendorf tubes and digested with 0.1% Collagenase type II from Cl. Histolyticum (Gibco) in Hanks Balanced Salt Solution without Mg^2+^/Ca^2+^ (Thermofisher) for 20-30 mins at 32°C with constant shaking. Once embryos were noticeably digested, cell solution was spun at 2000 g for 5 min at room temperature and the supernatant was removed. Cell pellet was resuspended with TrypLE Express (Thermofisher) and digested for 10 min at 32°C with constant shaking. Cell solution was inactivated with 10% Fetal Bovine Serum (Thermofisher) in Hanks Balanced Salt Solution without Mg^2+^/Ca^2+^ and filtered through a 70-μmcell strainer (Greiner Bio-One). Cells were pelleted at 2000g 5min room temperature and washed twice with 10% Fetal Bovine Serum (Thermofisher) in Hanks Balanced Salt Solution without Mg^2+^/Ca^2+^. Hoechst 34580 at a final concentration of 16.8 ug/mL was added to the cell solution and incubated for 30 mins at 28°C in the dark. Solution was then filtered through a 40-μmcell strainer (Greiner Bio-One), and propidium iodide was added at a final concentration of 5 ul/ml. FACS was performed on BD FACSInflux as described above, retaining only cells in G2/M phase based on Hoechst DNA content. Plates were processed for scDam&T-seq as described above.

### Immunofluorescent staining and confocal imaging of zebrafish embryos

Embryos at 15-somite stage were fixed in 4% PFA (Sigma) for 2 h at RT, followed by washes in PBS. Embryos were then washed three times in 4% sucrose/PBS and allowed to equilibrate in 30% sucrose/PBS at 4°C for 3-5 h. Embryos were suspended in Tissue Freezing Medium (Leica) orientated in the sagittal plane and frozen with dry ice. Blocks were sectioned at 8 μm and slides were rehydrated in PBS, treated with -20°C pre-cooled acetone for 7 min at -20°C, washed three times with PBS and digested with Proteinase K (Promega) at a final concentration of 10 ug/mL for 3 min, washed 1x PBS and incubated in blocking buffer (10% Fetal Bovine Serum, 1% DMSO, 0.1% Tween20 in PBS) for 30 min. Primary antibody was diluted in blocking buffer and slides incubated overnight at 4°C. Slides were washed the following day and incubated with the appropriate AlexaFluor secondary antibodies (1:500), DAPI (0.5 μg/mL) and Phalloidin-TRITC (1:200) diluted in blocking buffer for 1 h at RT. Slides were washed, covered with glass coverslips with ProLong Gold Antifade Mountant (Thermofisher) and imaged at 63X with a LSM900 confocal with AiryScan2 (Zeiss). Images were viewed and processed in Imaris 9.3 (Bitplane) and Adobe Creative Cloud (Adobe). Primary antibody: anti-H3K9me3 abcam ab8898 at 1:500 (Chandra et al., 2012).

### Processing DamID and scDam&T-seq data

Data generated by the DamID and scDam&T-seq protocols was largely processed with the workflow and scripts described in (Markodimitraki et al., 2020) (see also www.github.com/KindLab/scDamAndTools). The procedure is described in short below.

#### Demultiplexing

All reads are demultiplexed based on the barcode present at the start of R1 using a reference list of barcodes. In the case of scDam&T-seq data, the reference barcodes contain both DamID-specific and CELseq-specific barcodes and zero mismatches between the observed barcode and reference are allowed. In the case of the population DamID data, the reference barcodes only contain DamID-specific barcodes and one mismatch is allowed. The UMI information, also present at the start of R1, is appended to the read name.

#### DamID data processing

DamID reads are aligned using bowtie2 (v. 2.3.3.1) (Langmead & Salzberg, 2012) with the following parameters: “--seed 42 --very-sensitive -N 1”. For human samples, the hg19 reference genome is used; for mouse samples, the mm10 reference genome; and for zebrafish samples the GRCz11 reference genome. The resulting alignments are then converted to UMI-unique GATC counts by matching each alignment to known strand-specific GATC positions in the reference genome. Any reads that do not align to a known GATC position or have a mapping quality smaller than 10 are removed. In the case of bulk DamID samples, up to 64 unique UMIs are allowed per GATC position, while up to 4 unique UMIs are allowed for single-cell samples to account for the maximum number of alleles in G2. Finally, counts are binned at the desired resolution.

#### CELseq data processing

CELseq reads are aligned using tophat2 (v. 2.1.1) (Kim et al., 2013) with the following parameters: “--segment-length 22 --read-mismatches 4 --read-edit-dist 4 --min-anchor 6 --min-intron-length 25 --max-intron-length 25000 --no-novel-juncs --no-novel-indels --no-coverage-search --b2-very-sensitive --b2-N 1 --b2-gbar 200”. For mouse samples, the mm10 reference genome and the GRCm38 (v. 89) transcript models are used. For zebrafish samples, the GRCz11 reference genome and the adjusted transcript models published by the Lawson lab (Lawson et al., 2020) are used. Alignments are subsequently converted to transcript counts per gene with custom scripts that assign reads to genes similar to HTSeq’s (Anders et al., 2015) htseq-count with mode “intersection_strict”.

### Processing of ChIP-seq data

External ChIP-seq datasets were downloaded from the NCBI GEO repository and the ENCODE database (Davis et al., 2018). The external ChIP-seq data used in this manuscript consists of: H3K9ac ChIP-seq in mESC (ENCSR000CGP), H3K27me3 ChIP-seq in mESC (ENCSR059MBO), and H3K9me3 ChIP-seq in 6-hpf zebrafish embryos (Laue et al., 2019) (GSE113086). Internal and external ChIP-seq data were processed in an identical manner. First reads were aligned using bowtie2 (v. 2.3.3.1) with the following parameters: “-- seed 42 --very-sensitive -N 1”. Indexes for the alignments were then generated using “samtools index” and genome coverage tracks were computed using the “bamCoverage” utility from DeepTools (v. 3.3.2) (Ramírez et al., 2016) with the following parameters: “-- ignoreDuplicates --minMappingQuality 10”. For marks that exist in broad domains in the genome, domains were called using MUSIC (Harmanci et al., 2014) according to the suggested workflow (https://github.com/gersteinlab/MUSIC). For marks that form narrow peaks in the genome, peaks were called using MACS2 (v. 2.1.1.20160309) using the “macs2 callpeak” utility with the following parameters: “-q 0.05”.

### Computing the Information Content (IC) of DamID samples

The Information Content (IC) of a DamID sample is a measure of how much structure is in the detected methylation signal. It is essentially an adaptation of the RNA-seq normalization strategy called PoissonSeq (Li et al., 2012). Its goal is to compare the obtained signal to a background signal (the density of mappable GATCs), identify regions where the signal is similar to background, and finally compare the amount of total signal (i.e. total GATC counts) to the total signal in background regions. The IC is the ratio of total signal over background signal and can be used to filter out samples that contain little structure in their data.

As an input, we use the sample counts binned at 100-kb intervals, smoothened with a 250-kb gaussian kernel. The large bin size and smoothing are necessary when working with single - cell samples that have very sparse and peaky data and would otherwise be difficult to match to the background signal. As a control, we use the number of mappable GATCs in the same 100-kb bins, similarly smoothened. We subsequently remove all genomic bins that do not have any observed counts in the sample. Our starting data is then *X*, a matrix with size (*n*, *k*), where *n* is the number of genomic bins and *k* is the number of samples. Since we are comparing one experimental sample with the control, *k* is always 2. *X*_*ij*_denotes the number of counts observed in the *i*th bin of the *j*th sample. We first compute the expected number of counts for each *X*_*ij*_based on the marginal probabilities of observing counts in each bin and in each sample:

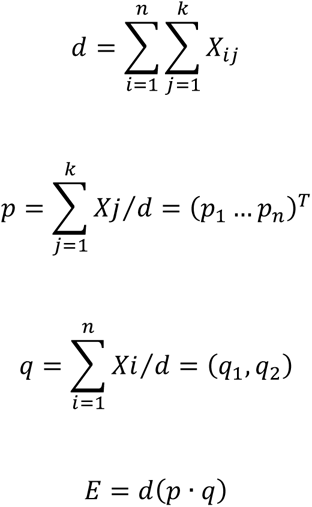

Where *d* is the total sum of *X*_*ij*_; *p*_*i*_is the marginal probability of observing counts in bin *i*; *p*_*j*_is the marginal probability of observing counts in sample *j*; and *E* is the matrix of size (*n*, *k*) where entry *E*_*ij*_is the expected number of counts in bin *i* for sample *j*, computed as *p*_*i*_*q*_*j*_*d*.

We subsequently compute the goodness of fit of our predictions compared to the actual counts per bin:

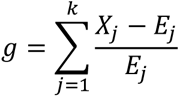

Where *g*_*i*_is the measure of how well the predictions of *E*_*i*_match the observed counts in *X*_*i*_in bin *i* . The better the prediction, the closer *g*_*i*_is to zero, indicating that the signal of the experimental sample closely resembles the background in bin *i*. Next, an iterative process is performed where in each step a subset of the original bins is chosen that exclude bins with extreme values of *g*. Specifically, all bins with a goodness of fit in the top and bottom 5^th^ percentiles are excluded to progressively move towards a stable set of bins where the sample resembles the background. After each iteration, the chosen bins are compared to the previous set of bins and when this has stabilized, or when the maximum number of iterations is reached, the procedure stops. In practice, convergence is usually reached after only a couple of iterations. The IC is then computed for the experimental sample as the ratio of its summed total counts to the sum of counts observed in the final subset of bins.

### Population DamID data filtering and analyses

The population DamID samples were filtered based on a depth threshold of 100,000 UMI-unique GATC counts and an IC of at least 1.1. Per Dam-construct, the best samples based on the IC were maintained. Samples were normalized for the total number of counts using reads per kilobase per million (RPKM). Normalization for Dam controls was performed by adding a pseudo count of 1, taking the per bin fold-change with Dam, and performing a log2-transformation, resulting in log2 observed-over-expected (log2OE) values. The UMAP presented in Figure 1B was computed by performing principal component analysis (PCA) on the RPKM-normalized samples (20-kb bins) and using the top components for UMAP computation in python with custom scripts. For the correlations presented in Figure 1C and S1C, the RPKM-normalized DamID values were normalized for the density of mappable GATCs and log-transformed. The Spearman’s rank correlation was then computed with the input-normalized ChIP-seq values of the various marks.

### Single-cell DamID data filtering and analyses

#### Filtering and normalizing scDamID data

Single-cell DamID samples were filtered based on a depth and an IC threshold. For the mouse samples, these thresholds were 3,000 unique GATCs and an IC within the range of 1.5 to 7 (the upper threshold removes samples with very sparse profiles); for zebrafish, these thresholds were 1,000 unique GATCs and an IC within the range of 1.2 to 7. For the zebrafish samples, chromosome 4 was excluded when determining depth and IC (and in all downstream analyses) since the reference assembly of this chromosome is poor and alignments unreliable. The quality of scDam&T-seq samples is determined separately for the DamID readout and the CEL-Seq2 readout. To preserve as much of the data as possible, we used all samples passing DamID thresholds for analyses that relied exclusively on the DamID readout. Wherever single-cell data was used, samples were normalized for their total number of GATCs, scaled by a factor 10,000, and log-transformed with a pseudo-count of 1, equivalent to the normalizations customarily performed for single-cell RNA-seq samples. To generate *in silico* populations based on single-cell samples, the binned UMI-unique counts of all single-cells were combined and normalization was performed equivalent to population DamID samples.

#### scDamID UMAPs

The UMAPs presented in Fig. 2A, Fig. 3C and Fig. 5C were computed by performing PCA on the depth-normalized single-cell samples and using the top components for UMAP computation. Since in EBs inactivation of chromosome X can coincides with a strong enrichment of H3K27me3/RING1B on that chromosome, we depth-normalized these samples using the total number of GATCs on somatic chromosomes. For the zebrafish samples, chromosome 4 was completely excluded from the analysis. For the mouse UMAPs, the single-cell data were binned at a resolution of 10-kb intervals, while for the zebrafish UMAPs, the resolution was 100 kb. Notably, when the first principal components showed a strong correlation to sample depth, it was excluded.

#### Single-cell count enrichment

Figures 2B-D show the enrichment of counts in ChIP-seq domains for all single-cell mESC samples. The count enrichment is equivalent to the more well-known Fraction Reads in Peaks (FRiP) metric, but has been normalized for the expected fraction of counts within the domains based on the total number of mappable GATCs covered by these domains. In other words, if the domains cover 50% of the mappable GATCs in the genome and we observe that 70% of a sample’s counts fall within these domains, the count enrichment is 0.7 / 0.5 = 1.4.

### Single-cell CELseq data filtering and analyses

#### Filtering CELseq data

Single-cell data sets were evaluated with respect to the number of unique transcripts, percentage mitochondrial reads, percentage ERCC-derived transcripts and the percentage of reads coming from unannoted gene models (starting with “AC” or “Gm”) and appropriate thresholds were chosen. For the EB data, the used thresholds were ≥1,000 UMI-unique transcripts, <7.5% mitochondrial transcripts, <1% ERCC-derived transcripts, and <5% transcripts derived from unannotated gene models. In addition, a small group of cells (29/6,554 ≍ 0.4%) from different time points, which formed a cluster that could not be annotated and was characterized by high expression of ribosomal genes, was removed from further analyses. For the zebrafish data, the used thresholds were ≥1,000 UMI-unique transcripts and <5% ERCC-derived transcripts. Only genes observed in at least 5 samples across the entire dataset were maintained in further analyses. The quality of scDam&T-seq samples is determined separately for the DamID readout and the CEL-Seq2 readout. To preserve as much of the data as possible, we used all samples passing CEL-Seq2 thresholds (independent of DamID quality) for transcriptome-based analyses.

#### Analysis of CELseq data with Seurat and Harmony

Single-cell transcription data was processed using Seurat (v3) (Stuart et al., 2019). First, samples were processed using the “NormalizeData”, “FindVariableFeatures”, “ScaleData”, and “RunPCA” commands with default parameters. Subsequently, batch effects relating to processing batch and plate were removed using Harmony (Korsunsky et al., 2019) using the “RunHarmony” command, using a theta=2 for the batch variable and theta=1 for the plate variable. Clustering and dimensionality reduction were subsequently performed with the “FindNeighbors”, “FindClusters” and “RunUMAP” commands. Differentially expressed genes per cluster were found using the “FindAllMarkers” command.

#### Integration with external single-cell datasets

The EB data was integrated with part of the single-cell mouse embryo atlas published by (Pijuan-Sala et al., 2019). The data was loaded directly into R via the provided R package “MouseGastrulationData”. One data set per time point was included (datasets 18, 14, 19, 16, 17, corresponding to embryonic stages E6.5, E7.0, E7.5, E8.0, E8.5, respectively). The atlas data and our own data was integrated using the SCTransform (Hafemeister & Satija, 2019) and the anchor-based intragration (Stuart et al., 2019) functionalities from Seurat. First, all data was normalized per batch using the “SCTransform” command. All data sets were then integrated using the “SelectIntegrationFeatures”, “PrepSCTIntegration”, “FindIntegrationAnchors”, and “IntegrateData”, as per Seurat documentation.

#### SCENIC

We used SCENIC (Aibar et al., 2017) on the command line according to the documentation provided for the python-based scalable version of the tool (pySCENIC) (van de Sande et al., 2020). Specifically, we ran “pyscenic grn” with the parameters “--method grnboost2”; “pyscenic ctx” with the parameters “--all_modules”; and “pyscenic aucell” with the default parameters. We used the transcription factor annotation and the transcription factor motifs (10 kb +/- of the TSS) provided with SCENIC. This yielded 414 activating regulons. We subsequently filtered regulons based on the expression of the regulon as a whole (at least 50% of cells having an AUCell score > 0 within at least one Seurat cluster) and based on the expression of the regulon transcription factor (detected in at least 5% of cells in at least one cluster) to retain only high confidence regulons. This resulted in 285 remaining activating regulons. However, repeating all analyses with the unfiltered set of regulons yielded the same trends and relationships.

### Linear Discriminant Analysis (LDA) classifier to assign samples to transcriptional clusters based on DamID signal

In both the EB results and the zebrafish results, we noticed that there was a substantial number of cells that passed DamID thresholds, but that had a poor CEL-Seq2 readout. Since most of our analyses rely on the separation of cells in transcriptional clusters (i.e. cell types) and cells with a poor CEL-Seq2 readout cannot be included in the clustering, these cells cannot be used in downstream DamID-based analyses. However, we noticed that the separation of different cell types was recapitulated to a considerable extent in low-dimensionality representations of the DamID readout (see the DamID-based UMAPs in Fig. 2A and Fig. 3D). Since cell-type information is captured in the DamID readout, we reasoned that a classifier could be trained based on cells with both good DamID and CEL-Seq2 readouts to assign cells with a poor CEL-Seq2 readout to transcriptional clusters based on their DamID readout.

To this end, we implemented a Linear Discriminant Analysis (LDA) classifier as described below.

#### Data input and preprocessing

As in input for the classifier, we used the binned DamID data of all samples passing DamID thresholds and the transcriptional cluster labels of these samples (samples with a poor CEL-Seq2 readout had the label “unknown”). The DamID data was depth-normalized (as described above) and genomic bins that contained fewer than 1 mappable GATC motif per kb were excluded, resulting in a matrix of size *N* x *M*, where *N* is the number of samples and *M* is the number of remaining genomic bins. For the EB data, a bin size of 10 kb was used, while a bin size of 100 kb was used for the zebrafish data. Subsequently, the pair-wise correlation was computed between all samples, resulting in a correlation matrix of size *N* x *N*. This transformation had two reasons: First, it served as a dimensionality reduction, since *N* << *M*. Second, it resulted in a data type that effectively describes the similarity of a sample with all other samples, including samples without a cluster label. Consequently, during the training phase, the classifier can indirectly use the information of these unlabeled samples to learn about the overall data structure. We found that using the correlation matrix (*N* x *N*) as an input for the classifier yielded much better results than using the original matrix (*N* x *M*).

To train the LDA classifier, we used two thirds (∼66%) of all samples with cluster labels (i.e. with a good CEL-Seq2 readout). Since the number of cells per cluster varied extensively, we randomly selected two thirds of the samples per cluster and thereby ensured that all clusters were represented in both training and testing. The training data thus consisted of the correlation matrix of size *N_train_* x *N* and a list of sample labels of size *N_train_*, where *N_train_* is the number of samples used for training. Consequently, we retained one third (∼33%) of labelled samples to test the performance of the LDA classifier, consisting of the correlation matrix of size *N_test_* x *N* and a list of sample labels of size *N_test_*, where *N_test_* is the number of samples used for testing. In summary, this split the samples into three groups: one group for training, one group for testing, and the group of unlabeled samples.

#### Training the classifier

For the implementation of the LDA classifier, we used the “LinearDiscriminantAnalysis” function provided in the Python (v. 3.8.10) scikit-learn toolkit (v. 0.24.2). The number of components was set to the number of transcriptional clusters minus one and the LDA classifier was trained using the training samples.

#### Testing the performance

To test the performance, the trained LDA classifier was used to predict the labels of the training set of samples. Predictions with a probability larger than 0.5 were maintained, while predictions with a lower probability were discarded (and the corresponding cells were thus not labelled). The predicted labels were subsequently compared to the known labels (Fig. S3E, S5.1C). In general, we found a very good performance for clusters with many cells, while the performance tended to be lower for clusters with few cells. This is as expected, since the number of samples for these clusters was also very low during training.

#### Predicting cluster labels for unlabeled samples

After establishing that the performance was satisfactory, the LDA was retrained, this time using all labelled samples. The actual performance on the unlabeled data is likely higher than the performance on the test data, since the number of samples used for the final training is notably higher. Finally, the cluster labels were predicted for the unlabeled samples. Once again, only predictions with a probability higher than 0.5 were maintained. Figure S3D shows the number of EB samples that were attributed to each cluster using the LDA classifier, as well as the number of samples that could not be attributed (“unassigned”).

### Defining PRC targets

First, we identified for each gene the region of 5 kb upstream and 3 kb downstream of the TSS. Only protein-coding genes and genes for non-coding RNA were considered. When the TSS domains of two genes overlapped, they were merged if the overlap was >4 kb, otherwise the two domains were split in the middle of the overlap. This resulted in 30,356 domains covering a total of 35,814 genes. Subsequently, for all single-cells, the number of observed GATC counts within each domain was determined. *In silico* populations per transcriptional cluster were generated by combining the counts of all cells belonging to each cluster per DamID construct. The *in silico* population counts were subsequently RPKM-normalized, using the total number of GATC counts on the somatic chromosomes of the combined single-cell samples as the depth (i.e. also counts outside the domains). Normalization for Dam controls was performed for the H3K27me3 and RING1B data per transcriptional cluster by adding a pseudo count of 1, taking the fold-change with Dam, and performing a log2-transformation, resulting in log2 observed-over-expected (log2OE) values. The correlation of the resulting H3K27me3 and RING1B values per cluster is shown in Figure S3F. We subsequently determined PRC targets as those genes that showed H3K27me3 and RING1B log2OE values >0.35 in at least one cluster. PRC targets were defined based on the *in silico* population of the H3K27me3 and RING1B data of the mESC cells (Fig. 2) and the EB clusters, excluding cluster 7. Cluster 7 was excluded, because it consisted of relatively few cells and the combined data was consequently sparse.

### ChromHMM of zebrafish *in silico* populations

In order to determine regions that were characterized by H3K9me3-enrichment in specific (sets of) cell types in the zebrafish embryo, we made use of ChromHMM (v. 1.22) (Ernst & Kellis, 2012, 2017). As input, we used the *in silico* H3K9me3 signal (log2OE) of all clusters that had at least 30 cells passing DamID thresholds for both Dam and MPHOSPH8 (clusters 0-11). The genome-wide signal at a resolution of 50 kb was used and the values were binarized based on a threshold of log2OE > 0.35. Bins that had fewer than 1 mappable GATC per kb were given a value of 2, indicating that the data was missing. As in all other analysis, chromosome 4 was excluded. The binarized values of clusters 0-11 were provided as input for the ChromHMM and the results were computed using the “LearnModel” function using the following parameters: -b 50000 -s 1 -pseudo. The number of ChromHMM states was varied from 2 to 10 and for each result the differences between the states (based on the emission probabilities) were inspected. We found that a ChromHMM model with 5 states was optimal, since this yielded the most diverse states and increasing the number of states just added redundant states with similar emission probabilities.

### Repeat enrichment in ChromHMM states

The RepeatMasker repeat annotations for GRCz11 were downloaded from the UCSC Genome Browser website (https://genome.ucsc.edu/). The enrichment of repeats within each ChromHMM state was computed either for repeat classes as a whole (Fig. S5.2A) or for individual types of repeats (Fig. 5I and S5.2C). To compute the enrichment of a repeat class/type in a ChromHMM state, the fraction of repeats belonging to that class/type that fell within the state was computed and normalized for the fraction of the genome covered by that state. In other words, if we observe that 70% of a certain repeat falls within state B and state B covers 7% of the genome, then the repeat enrichment is 0.7 / 0.07 = 10.

### GO term and PANTHER protein classs enrichment analysis

GO term and PANTHER (Mi et al., 2013) protein class enrichment analyses were performed via de Gene Ontology Consortium website (http://geneontology.org/). For Figure S4E, the list of PRC-regulated TFs was used as a query and the list of all TFs as a reference to determine enriched Biological Process GO terms. Only the top 10 most significant terms are shown. For Figure S5.1G, the list of genes in ChromHMM state A1 or B was used as a query and the list of genes in all ChromHMM states as a reference to determine enriched PANTHER protein classes. All hits are shown.

## Notes

### Competing Interest Statement

The authors have declared no competing interest.

